# Caveolin-1 dependent regulation of cell-matrix interphase in 3D collagen gels

**DOI:** 10.1101/2025.06.04.657900

**Authors:** Debasmita Mazumdar, Sujal Kataria, Gyanendra Prasad Panda, Atharva Kulkarni, Shivprasad Patil, Mamoni Dash, Nagaraj Balasubramanian

## Abstract

Cell and extracellular matrix (ECM) interactions are essential for maintaining tissue function and homeostasis. Changes in the biochemical or mechanical properties of the ECM can lead to diseases such as fibrosis or cancer. In a 3D microenvironment, cell-matrix interaction is vital to how cells sense and respond to biochemical and biophysical cues. This study examines the reciprocal interactions between fibroblasts and collagen in 3D hydrogels. We quantitatively measured changes in collagen branch number and junctions in 3D hydrogels using confocal reflectance microscopy and existing analysis protocols. This reveals the impact small changes in collagen concertation (1.0 vs 1.5 mg/ml) over time (15 minutes to 4 hours) have on 3D gels. Embedded in 3D hydrogels, wild-type mouse fibroblasts differentially affect collagen organisation in their immediate proximity with changing concentration and time. This regulation is interestingly lost in Caveolin-1 null fibroblasts with altered stiffness, mechanosensing and cytoskeletal regulation. Inhibition of the Rho-ROCK pathway (known to be changed in Caveolin-1 null fibroblasts) drives cellular protrusions and concentration-dependent 3D collagen organisation in wildtype fibroblasts, but surprisingly not in Caveolin-1 null fibroblasts. This depends on dynamin-dependent endocytosis, which, when inhibited, disrupts ROCK-dependent protrusions and alters collagen organisation in 3D collagen. Together, these observations quantitatively demonstrate how cells respond at the cell-matrix interphase to subtle changes in collagen concentration and organisation in 3D hydrogels, regulated by the presence of Caveolin-1.

**Statement of Significance:** This study highlights the critical role of Caveolin-1 in cell-mediated regulation of the extracellular matrix (ECM) in a 3D microenvironment. Using confocal reflectance microscopy, we demonstrate that subtle changes in collagen concentration significantly influence collagen organisation over time. While wild-type fibroblasts exhibit dynamic, concentration-dependent regulation of the cell-matrix interphase, Caveolin-1 null fibroblasts lose this regulation. Furthermore, Rho-ROCK pathway inhibition promotes protrusions and collagen reorganisation in wild-type cells but not Caveolin-1 null fibroblasts. Dynamin-dependent endocytosis also emerges as a key modulator of this process. These findings provide quantitative insights into how mechanosensing and cytoskeletal regulation, dependent on endocytosis can control the cell-matrix interphase, with possible implications for understanding disease-associated matrix changes.

## Introduction

The extracellular matrix (ECM) comprises various proteins such as collagen, fibronectin, and laminin along with proteoglycans and glycosaminoglycans, as well as enzymes, proteases, and their inhibitors that impact cells through their biochemical and biophysical cues (1–3). These proteins and their matrix properties can vary significantly between different tissue types. Cells in the tissue play a crucial role in the making and maintaining the ECM through development. They also remodel the matrix in response to injury or disease to facilitate adaptation and repair. The reciprocal interaction between cells and their surrounding ECM is a dynamic process that depends not only on biochemical factors but also on the biophysical properties of the matrix. In diseases like cancer, changes in both the cells and the ECM contribute to disease progression.

Cells sense mechanical changes in their surrounding microenvironment, rapidly converting them into biochemical signals that ultimately trigger a biological response. This process is known as mechanotransduction (4,5). This response can be intracellular, such as changes in protein expression or function, or extracellular, involving the secretion, degradation, or remodelling of the ECM (6). Vital membrane mechanosensory proteins, such as integrins and caveolins, play a pivotal role in cellular mechanotransduction (7,8). They do so by diverse means.

Caveolin-1 (Cav-1), a crucial membrane-attached mechanosensory protein, is primarily located in caveolae, a specialised cholesterol-rich plasma membrane invagination with a characteristic omega shape (∼80 nm in size) structure. Caveolae functions as a mechanosensor, partly as a membrane reservoir and an endocytic pathway in cells (8–10). Cav-1 is also present in focal adhesions (FAs), which could help sense and respond cues from the ECM (10). In addition, Cav-1 influences ECM organisation by regulating its alignment and stiffening at the cell surface through its modulation of actin polymerisation (11). This dynamic interplay between Cav-1 and the ECM affects cellular homeostasis and tissue function.

Cells regulate the ECM through biochemical and biophysical means. Biochemically, the secretion of ECM proteins, matrix metalloproteinases (MMPs), or their inhibitors can all modulate matrix composition and organisation. Biophysically, cells modify the ECM by protrusive, contractile, or volumetric forces (5,12). The cellular actin cytoskeleton is vital for force generation and matrix remodelling. The polymerisation and dynamics of actin filaments are tightly controlled by Rho family small GTPases, such as RhoA, Rho-associated kinase (ROCK), Rac1, and Cdc42. Cav-1 is also seen to inhibit MMP secretion while promoting ROCK signaling and actin contractility (13,14).

In tumours, the ECM undergoes significant changes. These alterations are primarily driven by increased ECM deposition, mainly collagen, by transformed cells, along with differential crosslinking and alignment of these proteins (15). As a result, the tumour ECM is stiffer and denser than normal tissue. These changes in the mechanical characteristics of the ECM play a crucial role in regulating the behaviour of cancer cells, including their migration and metastasis (16). This also highlights the importance of maintaining ECM homeostasis, in its composition and mechanical properties in tissues.

Collagen accounts for approximately 30% of the total protein mass in the body and up to 90% of the ECM in various tissues (17). This fibrous protein is secreted by cells and consists of three α chains that combine to form a triple-helix structure known as procollagen. Procollagen assembles into fibrils, which then polymerise to form collagen fibres. They crosslink with each other to create the structural meshwork that provides mechano-physical support to cells (18,19). Various factors, including collagen concentration, the pH of the microenvironment, ionic strength, and temperature, tightly regulate the polymerisation and crosslinking of collagen. These factors influence basic properties of collagen fibres, such as their length, thickness, crosslinking density, and pore size. Even small changes in collagen concentration can alter these parameters, potentially disrupting their mechanical homeostasis and affecting cell behaviour (18,20,21). This also emphasises the importance of cells being able to sense and respond to subtle variations in their matrix microenvironment to eventually regulate cellular function.

In the 3D matrix, the cell in being surrounded by the ECM, creates a dynamic interface for reciprocal regulation. Cell-ECM interactions are mediated by membrane receptors like integrins, which facilitate signalling and physically link the intracellular actin cortex to the ECM network. This biochemical and mechanical coupling reinforces each other, shaping cellular behaviour(22). These emphasise the need to evaluate the cell-matrix interaction and interphase in 3D, to understand their impact on cells in physiology better.

In this study, we investigated whether fibroblasts can detect subtle variations in collagen concentrations in their 3D microenvironment and how they respond to these differences. A previous study from our lab developed a protocol that allows a detailed examination of the properties of polymerised collagen within gels using Z-stack reflectance images (23). This approach enables us to observe characteristics such as branch junctions, total branch count, branch length, and more, providing insights into the nature of collagen polymerisation and how the presence of cells can impact it. The organisational nature of this cell-matrix interphase, its regulation by cell stiffness, cytoskeletal organisation, and the role of endocytosis were evaluated.

## Materials and Methods

### Materials

#### Cell culture

Cell culture media DMEM was purchased from Thermo Fisher-12800017. FBS (10270106) and Pen-Strep (15140122) were ordered from Gibco, Thermo Fisher.

#### For collagen gel preparation

Rat tail collagen 1 (3 mg/ml) was purchased from Gibco, Thermo Fisher Scientific (A1048301). 10X DPBS (Gibco, Thermo Fisher Scientific-14200075), 1X DPBS (Gibco, Thermo Fisher Scientific-14190136), and cell culture grade water (Gibco, Thermo Fisher Scientific 15230001) were purchased from Gibco. 1N NaOH was purchased from hi media (TCL002). The gels were prepared in an 8-well glass bottom lab-tek chamber (Nunc-155409).

#### Reagents

Phalloidin conjugated with Alexa Fluor 488 (A12379) and Alexa Fluor 594 (A12381) were purchased from Invitrogen, Thermo Fisher. Cholera Toxin subunit B conjugated with Alexa Fluor 488 (C22841) was purchased from Invitrogen Thermo Fisher Scientific. Transferrin. Latrunculin A (L5163), Y 27632 (Y0503) and Dynasore hydrate (D7693) were purchased from Sigma.

#### Antibody

Caveolin-1 HRP Rabbit Polyclonal antibody from Santacruz SC894 was used for western blot, and Caveolin-1 rabbit monoclonal antibody from cell signalling 3267S has been used for immunofluorescence.

#### Plasmids

Caveolin-1 mCherry was obtained from Dr Miguel Del Pozo (Vector DB no. 3553). Dynamin-K44A-GFP was purchased from Addgene 22301. LifeAct GFP was borrowed from Dr Aurnab Ghosh’s lab.

### Methods

#### Cell Culture

Wild Type Mouse Embryonic Fibroblasts (WT MEFs) and Caveolin-1 Null Mouse Embryonic Fibroblasts (Cav-1 Null MEFs) cell lines were a kind gift from Dr Richard Anderson’s lab (University of Texas Health Sciences Center). The cells were cultured in high-glucose Dulbecco’s Modified Eagle’s Medium supplemented with 5% Fetal Bovine Serum and Pen-Strep (Thermo Fisher) at 37°C in a 5% CO_2_ incubator. Cells were checked regularly for bacterial and mycoplasma contamination.

Cells were transfected with DNA using lipofectamine 2000 reagent (Invitrogen) for 14 hours, after which media was changed to regular 5% DMEM and incubated for 24 hours before using for experiments. The transfection efficiency was checked after 24 hours and before the experiment.

#### 3D collagen gel preparation

1.0 and 1.5 mg/ml 3D collagen gels were prepared with rat tail collagen I (3 mg/ml), 10X DPBS, distilled water, cells suspended in 1X PBS and 1N NaOH (to bring up the pH to 7.4) in a pre-decided ratio (Protocol adopted from Gibco (24)) and added to a well of 8-well glass bottom Lab Tek chamber. They were incubated at 37°C in a 5% CO_2_ incubator for 30 minutes. After polymerisation, 5% DMEM was added to the gels and incubated for 15 mins or 4 hours as needed. Gels were fixed or imaged live as needed. The same protocol was used to make gels without cells.

#### Latrunculin A, Y27632 and Dynasore Treatment

For the treatment with latrunculin A (Lat A), Y27632 or Dynasore, WT MEFs and Cav-1 Null MEFs were seeded on coverslips and incubated for 12 hours and treated with drug at required concentrations for 2 hours. Controls were treated with the same volume of DMSO. Coverslips were fixed with 3.5 % PFA for 15 min at room temperature (RT) and stained with phalloidin (1: 500 in 1X PBS) for 1 hour at RT, mounted using fluoramount mounting medium (Southern Biotech, Cat. #0100-01). For 3D collagen gels, cells were similarly treated. After 2 hours, drug or DMSO at the right concentration was added to collagen solution with cells their volume adjusted for polymerisation. 5% DMEM with drug or DMSO was added to gels for required incubation time.

#### Fixation and phalloidin staining of the cells embedded in the 3D collagen gels

Collagen hydrogels, with embedded cells are fixed with 4% PFA made with 5% sucrose for 30 min at RT, followed by three 1X PBS washes and incubated with Alexa Fluor 488/594 phalloidin diluted in 1X PBS (1:300) at 4°C overnight. Gels were washed with 1X PBS next day and stored at 4°C until imaged.

#### Immunofluorescence staining for Caveolin-1

Cells seeded on glass coverslips with or without Poly-L-Lysine (PLL) coating were fixed using 3.5% PFA for 15 min at RT. Permeabilised with 0.05% TritonX in PBS for 10 min at RT, blocked using 5% BSA in 1X PBS for 1 hour at RT. Incubated with Anti-Caveolin-1 antibody (1:200 dilution in 5% BSA) for 2 hours at RT, washed with PBS and incubated with fluorescent-tagged secondary antibody (1:1000 dilution in 5% BSA) for 1 hour at RT. The coverslips were washed with PBS and mounted using Fluoramount.

For cells embedded in 3D collagen gels we adapted a known protocol (25). After fixation, gels were permeabilised using 0.5% TritonX for half an hour, washed thrice with PBS, and incubated with primary antibody (1:100 dilution in 1% BSA) overnight at 4°C. Next day they were incubated with fluorescent-tagged secondary antibody (1:500 dilution in 1% BSA) overnight at 4°C. Gels were washed with PBS and stored at 4°C until imaged.

#### Imaging of collagen gels by confocal microscopy

Collagen gels with or without cells after fixation, were imaged as a Z stack using a Zeiss 780 multiphoton microscope. Collagen was imaged using reflectance mode with 488 laser and MBS T80/R20 reflection filter, and cells using confocal mode with the required laser. Z stack images were taken at 63X magnification and 2X digital zoom (keeping the cell at the centre), pinhole 1 AU with a step size of 0.32/0.34 um. All the images were analysed in ImageJ.

#### Live Imaging of cells in 3D collagen

For live imaging of cells in 3D collagen gels, cells were transfected with LifeAct GFP and embedded in gels as described earlier. After the required incubation time, they were imaged in a temperature-controlled chamber set to 37°C. Cross-section confocal and reflectance images were taken at 11-second intervals for 50 frames.

#### Analysis of collagen organisation

The Z stack collagen gel images were binarised for analysis using a macro created in ImageJ (23). These binarised images were skeletonised using the “Skeletonise 2D/3D” plugin, were further run through the “Analyse Skeleton 2D/3D” plugin to measure the parameters of the polymerised collagen in the given image (shown in Figure 1A). This is normalised by the volume of the images calculated from the summation of the area of all the XY planes multiplied by the step size.

**Figure 1:**
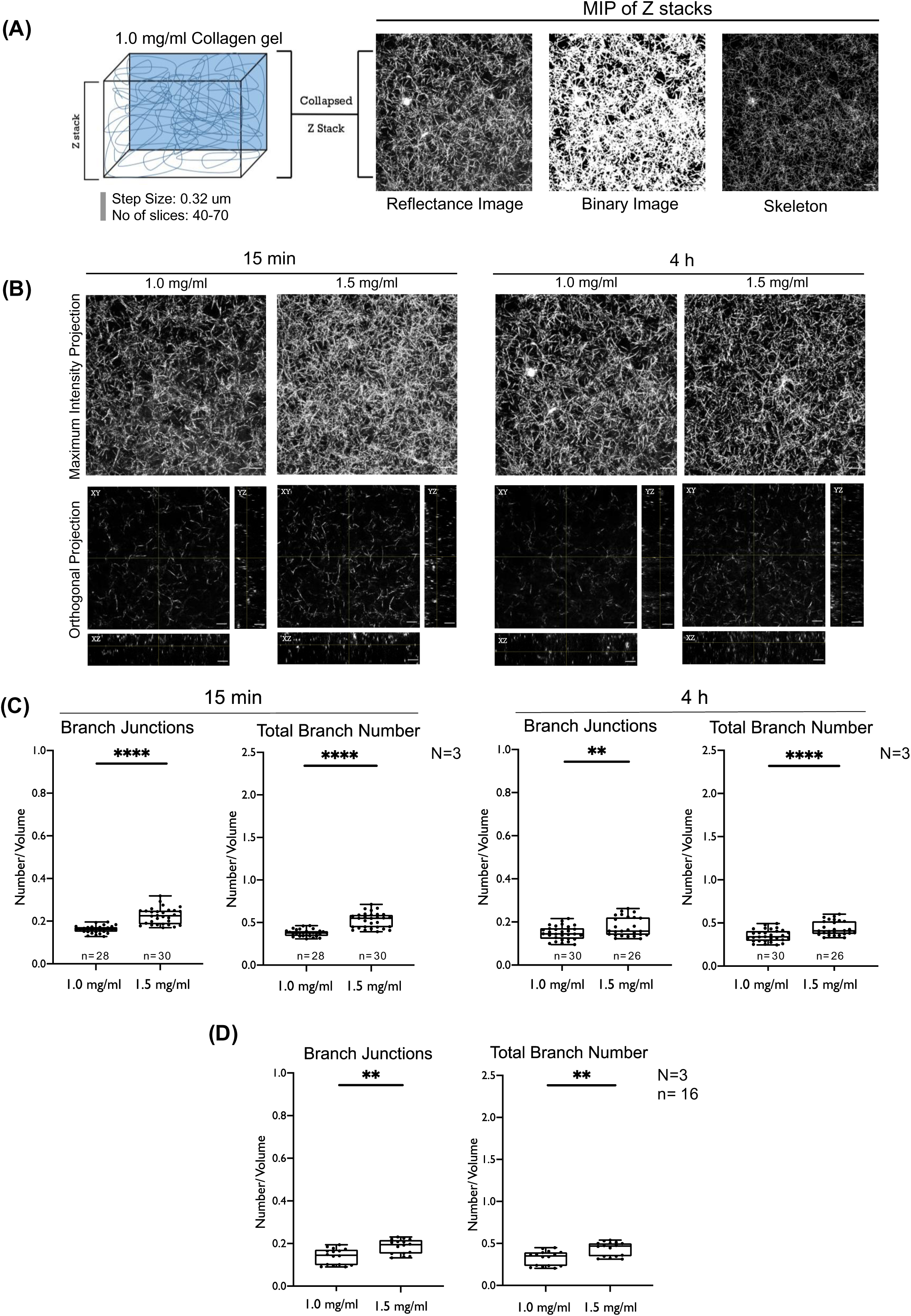
Collagen organisation (branch junctions and total branch number) in 3D collagen hydrogels at two different concentrations at different time points. **(A)** Image analysis pipeline for 3D collagen gel image using Image J. Representative cartoon of Z stack reflectance images; collapsed, binarised and skeletonised. Representative maximum intensity projection (MIP) images for 1.0 mg/ml collagen shown. **(B)** Representative MIP (top) and orthogonal projection (OP) (bottom) in XY, XZ and YZ plane for 3D collagen gels at 1.0 mg/ml and 1.5 mg/ml at 15 min or 4 hours. Box plot of Branch Junctions and Total Branch Number of Collagen gels of 1.0 mg/ml and 1.5 mg/ml concentrations at **(C)** 15 min and 4 hours with fixation (n= 26 to 30 Z-stack images from N=3 independent experiments) and **(D)** 1.0 mg/ml and 1.5 mg/ml collagen gels without fixation (n=16, N=3). Graphs show all data points with median and quarters. Error bars map the spread of data points. The number of junctions and branches is normalised to the volume of the image (Y axis). Scale Bar: 5 μm. Statistical analysis: Unpaired T test (**P<0.001, ****P<0.00001)

To analyse collagen organisation near the cell, a 50-pixel ROI around the cell was made. Phalloidin-stained cell in the Z stack image was used to create a mask of the cell. The cell mask was generated for each plane of the Z stack for each cell. Then, the ROI of the mask was put into the binarised reflectance channel (as explained earlier) of the respective image. Signal inside the ROI was used to remove the reflection of the cell. The ROI was increased by 50 pixels using the “Enlarge” command and signals outside the enlarged ROI were removed from the “cell removed reflectance Z stack image”. The image left is of collagen signal in a 50-pixel region representing the immediate surrounding of the cell **(Fig 2A)**. This new binary collagen image is skeletonised and analysed. The volume of this ROI through the Z stack is calculated from the area of the cell and the 50-pixel increased ROI. The area in this ROI was measured throughout the Z stack, and the sum of it was multiplied by the step size, to give us its volume. This was followed by subtracting the cell volume from the 50-pixel increased ROI, which gives us the volume of the 50-pixel region, used to normalise the data. The 50-pixel ROI was moved to a corner of the image away from the cell (X=-250, Y=250 pixels), and the collagen organisation was measured, as explained earlier. This protocol adapts the earlier published method from the lab (23).

**Figure 2:**
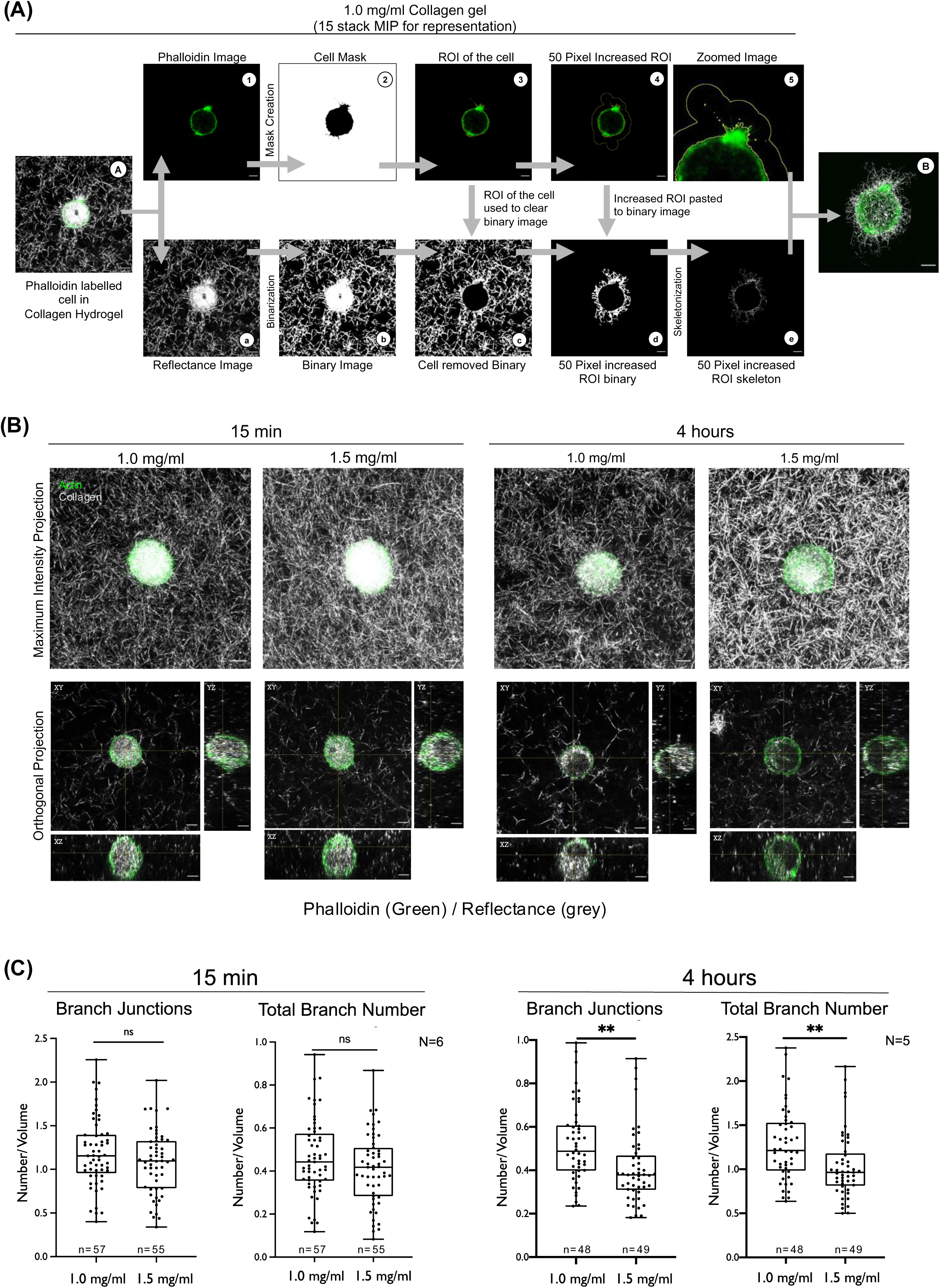
Collagen organisation in 3D gels near WT MEFs. **(A)** Collagen organisation analysis steps shown in flow chart marked by arrows. Representative MIP of 15 cross-section WT MEF Phalloidin Image and Reflectance Image are shown. Their combined image (A) is split as (1) a Phalloidin image (Upper panel), (2) a cell mask was created, (3) the ROI of the mask made, and (4) increased by 50 pixels, (5) zoomed image for 50 pixels increased ROI made. The (a) reflectance image (Lower panel) is,(b) binarised, (c) signal from the cell removed using the cell ROI (upper panel: 3) to create the cell removed binary image, (d) region outside the increased ROI (upper panel: 4) cleared to make the “50 Pixel increased ROI” binary image, (e) this image is then skeletonised. A combined image of a cell labelled with Phalloidin 488 with the skeletonised collagen fibers shown at the end. **(B)** Representative MIP and OP (in XY, XZ and YZ plane) images for WT MEF embedded collagen gels at 1.0 mg/ml and 1.5 mg/ml after 15 min and 4 hours, labelled green with Phalloidin (actin) and grey (reflectance). **(C)** Graphs show all data points with median and quarters on both sides for Branch Junctions and Total Branch Number (normalised to the volume of ROI) for collagen near WT MEFs at 1.0 mg/ml and 1.5 mg/ml collagen gels at 15 min (n= 57, n=55, N=6) and 4 hours (n=48,n=49, N=5). Error bars map the spread of data points. Scale Bar: 5 μm. Statistical analysis: Unpaired T test (**P<0.001).

#### Cell surface labelling and endocytosis

WT MEFs embedded in collagen gels, after 3 hours of incubation, were labelled with 2 ug/ml CTxB 488 and 25 ug/ml Transferrin 561 on ice for 30 minutes. Once labelled, the excess reagent was washed with PBS and cells incubated at 37°C for 60 min. This was done with and without drug when needed.

#### Atomic Force Microscopy (AFM)

The nanoindentation experiments on the cells were performed using JPK Nanowizard II AFM. The soft tip-less cantilevers of stiffness ∼0.1 N/m from MikroMasch (model-HQ: CSC38/tipless/Cr-Au) were used by attaching a 5 µm (diameter) glass bead using the lift-off method described elsewhere (26). Before each experiment, both the photodetector sensitivity and the cantilever’s force constant were determined. The detection sensitivity, measured in nanometers per volt (nm/V), was calculated by analyzing the slope of the approach curve obtained at the deep contact region between the cantilever and the glass slide. The force constant of the cantilever was evaluated using the thermal tuning method integrated within the JPK software.

For the AFM studies, 50,000 WT MEFs or Cav-1 Null MEFs were seeded on glass coverslips, incubated for 12 hours and AFM measurements done on live cells. Cells adherent for 12 hours, were treated with Lat A, Y27632 or Dynasore for 2 hours, and the stiffness was measured. Cells seeded on coverslips coated overnight at 4°C with 10 ug/ml PLL diluted in 1X PBS were incubated for 30 min and used for AFM measurements. Cells grown and processed in tissue culture hoods were taken to the AFM lab for measurements.

Force-deformation experiments were conducted on individual cells. The measurements were performed at a speed of approximately 2 µm/s over a 1 × 1 µm² area, employing a 7 × 7 grid. A sampling rate of 2 kHz was maintained throughout the experiments. The experimentally recorded raw parameters were the cantilever deflection (d_cant_, in Volts) and the base piezo displacement (d_pz_, in Volts), where the piezo displacement corresponds to the movement of the cantilever base. Both d_cant_ and d_pz_ were converted into meters by multiplying them with photodetector and base-piezo sensitivity, respectively.

The force-deformation curves are fitted using the Hertz model for a spherical bead of 2.5 um radius pressed against a flat surface, and the Poisson ratio of cells is assumed to be 0.5. The force (F) exerted on the cells was determined by multiplying the cantilever deflection (de_cant_) by the cantilever’s stiffness, and the deformation of the cell was obtained by subtracting the cantilever deflection from the base-piezo displacement.

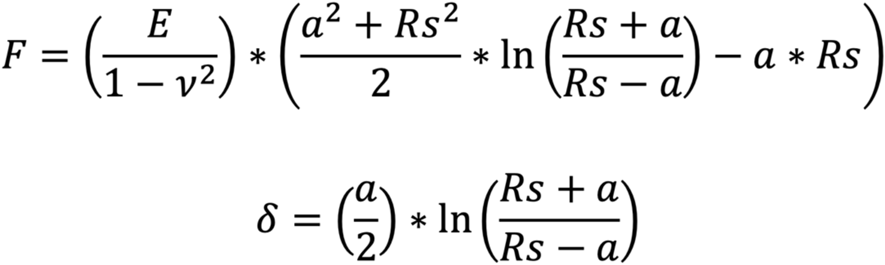

Here, F is measured by the cantilever possessing the bead, which is pressed against the cell. R_s_ is the bead radius, δ is the deformation in the cell, E is Young’s modulus, v is the Poisson ratio, and a is the contact radius. This analysis uses JPK’s data processing software to obtain Young’s modulus (E) of cells.

#### Rheology Measurements

Collagen hydrogels were prepared at 1.0 and 1.5 mg/ml concentrations with and without cells (WT MEFs and Cav-1 Null MEFs), 15 min or 4 hours after polymerisation using a rheometer (MCR 102E, Anton Paar). The hydrogels’ elasticity, crosslinking, and mechanical strength were evaluated using frequency sweep measurements performed on a modular compact rheometer (MCR 102E, Anton Paar). The analysis was conducted using aluminium plates in a parallel plate geometry with a plate diameter of 25 mm. For the frequency sweep analysis, the frequency range was varied from 10 to 0.1 rad/s while maintaining a constant shear strain of 0.1%. All measurements were carried out at 37°C. The data were processed using RheoPlus software, and key parameters such as the storage modulus (G’) and loss modulus (G”) were extracted from the processed data.

#### Statistical analysis

All statistical analyses were done using the Unpaired T-Test or Mann-Whitney test (to compare two groups of data) or one-way ANOVA (to make the comparison between more than two groups of data) in Prism Graphpad analysis software.

## Results

### Quantitative evaluation of fibrillar organisation in 3D collagen gels, dependent on concentration and incubation times

Subtle differences in collagen concentration in 3D gels could affect its organisation and changes over time could be vital to the behaviour of cells in these conditions. Local variations in extracellular matrix occur in *in vivo* tissue scaffolds in disease conditions (27–30). We hence evaluated 3D collagen gels made with rat tail collagen at a 1.0 mg/ml (LOW) and 1.5 mg/ml (HIGH) concentration at early (15 min) and late (4 hours) time points of incubation. These hydrogels were prepared as described in the methods, allowing collagen to polymerise into fibers, which crosslink with each other to give rise to a defined 3D organisation. These were analysed using an image J pipeline described earlier (23) **(Fig 1A, 1B)**. This evaluated fiber junctions, representing collagen crosslinking and the total fiber number in gels when normalised to gel volume. Collagen branch junctions and branch numbers are significantly higher in 1.5 mg/ml collagen than 1.0 mg/ml after 15 min incubation **(Fig 1C)**. After four hours of incubation, these differences were reduced but remained significant **(Fig 1C)**. When compared over time (15 min and 4 hours) 1.5 mg/ml gels showed a significant drop in branch junction (∼21%) and numbers (∼17%). No significant change was seen for 1.0 mg/ml hydrogels over time. This could reflect possible time-dependent relaxation of the gel, as has been reported earlier in the literature (27,29,30). Burla et al. have also suggested that the relaxation rate depends on the connectivity and plasticity of collagen gels (28) which could further depend on collagen concentration. This explains differences in relaxation rates and, hence, the change in collagen organisation observed in 1.5 mg/ml gels relative to 1.0 mg/ml.

To nullify the possibility of the artefact of paraformaldehyde fixation, “live” collagen gels (without fixation) were also evaluated. The collagen organisation showed comparable branch junctions and numbers and differences in the organisation between concentrations, as seen earlier for fixed gels **(Fig 1D)**. This supports using this imaging and analysis protocol for 3D collagen hydrogels to evaluate their behaviour with cells and study small changes (if any) in their organisation across collagen concentrations and time. Analysis of reflectance-imaged collagen gels without needing collagen to be stained could also provide an additional advantage of evaluating its organisation using this protocol.

### Effect embedded fibroblasts have on collagen organisation in 3D gels over time

Cells embedded in 3D collagen hydrogels interact with the matrix via integrins that can modulate collagen fibre organisation, alignment, and orientation in the immediate surroundings. The extent to which a single cell affects collagen organisation in its immediate surroundings and the cellular pathways that mediate this remains worth testing. In this study, WT MEFs were added to collagen, followed by its polymerisation, allowing cells to be embedded in the 3D matrix. Cell numbers were kept low to allow for single-cell imaging. The cell membrane was labelled with cholera toxin-B (CTxB), which binds GM1 and actin labelled with phalloidin, marking the cortex. In 3D gels, they closely overlap in untreated cells (Sup Fig 1.A). Using actin-labelled cells, the collagen gel near the cell (the cell boundary) was defined, and a 50-pixel region of interest (ROI) (3.25 µm) around the cell. This represents an extracellular region representing the cell-matrix interphase (Fig 2A). GM1 labelling was used under conditions where actin is disrupted as discussed later.

The 3D collagen reflectance signal from this narrow 50-pixel region surrounding the cell was analysed and compared. This collagen Z-stack image, when skeletonised, revealed a prominent collagen region with increased branching adjacent to the cell membrane. This could result from a more significant collagen accumulation, as reported earlier(31,32). This region is visible 15 min and 4 hours post-polymerization (Sup Fig 1B). For ease of understanding, we refer to this region as the “collagen cortex”.

Collagen organisation, when measured in this region and normalised to the ROI volume **(Fig 2B)** revealed WT MEFs to distinctly impact collagen organisation at both 1.0 and 1.5 mg/ml concentrations near the cell (18,32). At early 15 min, in the presence of cells, the number of branches and junctions in the immediate cell vicinity significantly increased compared to gels without cells **(Fig 2C)**, corroborating the presence of a visible collagen cortex. This increase persists at the late 4-hour time point **(Fig 2C)**. At 15 min, along with this, the number of branches and junctions in the immediate cell vicinity were not significantly different between the two collagen concentrations (1.5 mg/ml interestingly showing a trend toward being lower than 1.0 mg/ml) **(Fig 2C)**. After 4 hours, this trend becomes a significant decrease in branch junctions and numbers in the 1.5 mg/ml gel compared to 1.0 mg/ml **(Fig 2C).**

To further validate these changes, we used the 50-pixel cell periphery ROIs to compare the collagen organisation at regions away from the cells. ROIs were moved from the cell vicinity by adjusting their X and Y by 250 pixels and analysed **(Sup Fig. 1C, 1D)**. As an additional control, 50-pixel ROIs were also pasted into images of gels without any cells and analysed, as shown earlier **(Sup Fig. 1E, 1F).** Collagen branch junctions and numbers in ROIs away from the cell **(Sup Fig. 1C, 1D)** were comparable and similar to those in gels without cells **(Fig 2C, 2D)**. At 15 min, the 1.5 mg/ml gel had more branches and junctions than the 1.0 mg/ml gel. After 4 hours, this difference drops distinctly **(Sup Fig. 1D, 1F),** as was also seen for gels **(Fig 1C)**. This confirms the observed changes in collagen organisation at regions of the immediate vicinity of cells are indeed due to the impact of cell-matrix interactions in 3D gels.

Studies by Malandrino et al. have shown that in a 3D microenvironment, cell-ECM interactions are dependent on cell filopodia-mediated dynamic interactions. This drives time-dependent collagen accumulation in the immediate surroundings of the cell. This inversely correlates with ECM crosslinking (31,33), suggesting matrix concentrations with low crosslinking could accumulate in higher amounts. The significant increase in branches and junctions could reflect cross-linking changes. Expectedly, 1.0 mg/ml accumulates around the cell more, reflected as an increase in branch number and junctions over 1.5 mg/ml. That these differences are seen on small changes in collagen concentration could further reflect how wild-type mouse fibroblasts affect 3D gels.

### Altered mechanosensing and signalling in Caveolin-1 null fibroblast. Effect on collagen organisation

Much of how WT MEFs behave and affect 3D collagen gels in their immediate vicinity could be mediated by their mechanical properties, mechano-responsiveness and resulting signalling (8,11,35–37). Cav-1 knockout fibroblasts affect matrix stiffness-dependent cell spreading and signalling (34). 3D collagen gels with embedded Cav-1 Null MEFs have been shown to have significantly lower contractibility than WT MEFs (11,35). All these suggest that WT MEFs and Cav-1 Null MEFs with differing mechano-responsiveness could affect their surrounding microenvironment differently as well.

We hence compared the behaviour of WT and Cav-1 Null MEFs in 3D collagen gels. The loss of Cav-1 in Cav-1 Null MEFs was confirmed by western blot and immunostaining **(Fig 3A)**. The stiffness of WT MEFs and Cav-1 Null MEFs when adherent and spread on glass was tested using AFM. This showed WT MEFs to be up to two times stiffer than Cav-1 Null MEFs **(Fig 3B)**. This stiffness of cells is, in part, a characteristic of the cells and their response to external stimuli. The composition and structure of the cell membrane and the cytoskeletal organisation and dynamics are major regulators of cell stiffness (36,37). Both of these are affected by the loss of caveolin-1 and, hence caveolae (8).

**Figure 3:**
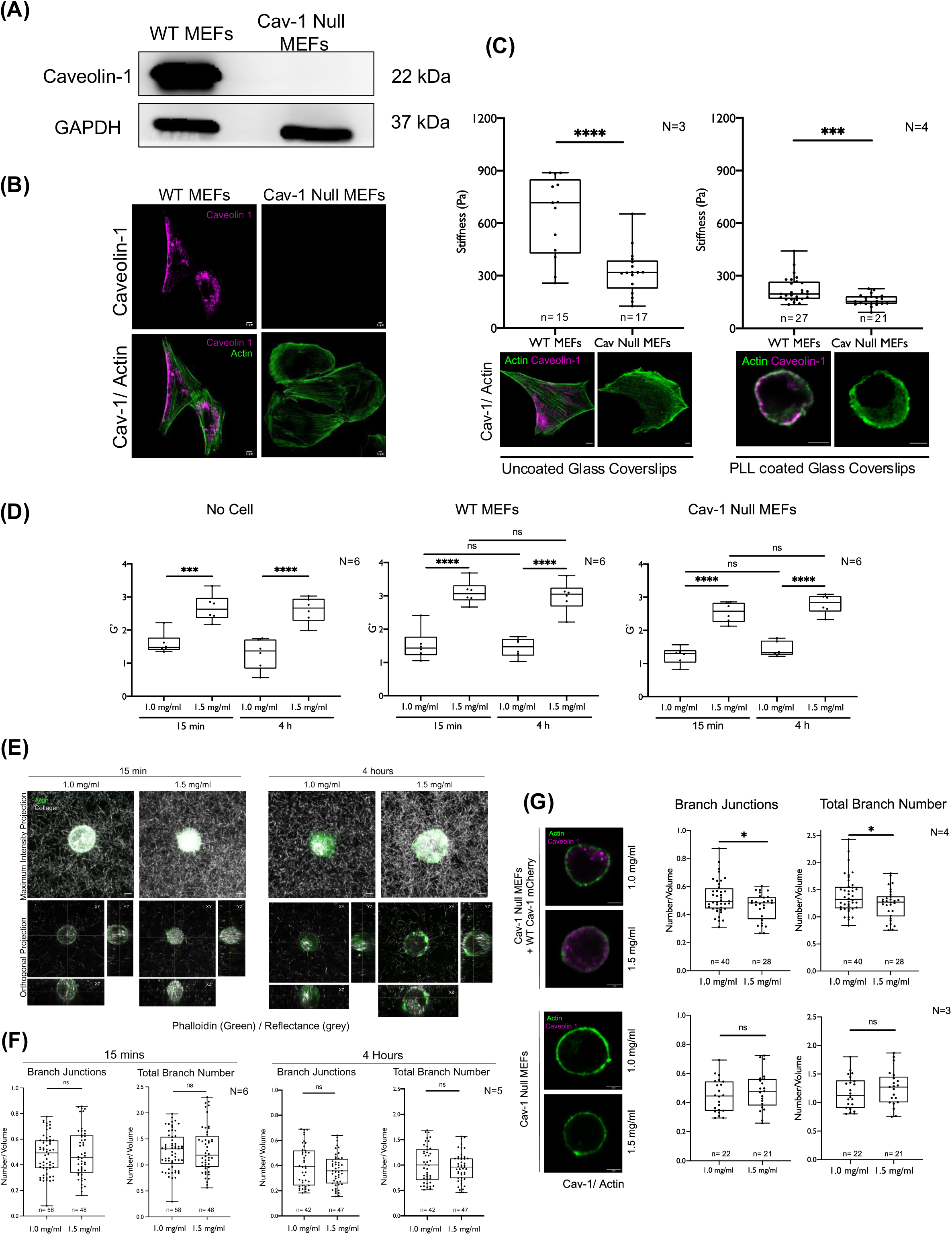
Cav-1 Null MEFs regulate collagen organisation differently than WT MEFs: **(A)** Western Blot detection of Caveolin-1 (22 kDa) and GAPDH (37 kDa) in WT MEFs and Cav-1 Null MEFs, **(B)** immunostained for Cav-1 (magenta) in cells adherent on 2D glass; stained with Phalloidin (green). **(C)** Stiffness (Pa) of WT MEFs and Cav-1 Null MEFs on uncoated (n=15,17, N=3) and Poly-L-Lysine (PLL) coated (n=27,21, N=4) glass coverslips measured using Atomic Force Microscopy (AFM). Representative images show both cells immunostained for Cav-1 (magenta) and actin (green). **(D)** Stiffness of 1.0 mg/ml and 1.5 mg/ml collagen gels measured using parallel plate rheology. Graphs represent stiffness (G’) of 1.0 mg/ml and 1.5 mg/ml collagen gels with no cell, with WT MEFs or Cav-1 Null MEFs at 15 min and 4 hours after polymerisation (N=6). **(E)** Z stack images of collagen gels with Cav-1 Null MEFs in 1.0 mg/ml and 1.5 mg/ml concentrations at 15 min and 4 hours. Representative MIP and OP (in XY, XZ and YZ plane) images, cells labelled with Phalloidin (green-actin) and reflectance channel (grey-collagen). **(F)** Graphs show branch junctions and total branch number of collagen in 50 pixel ROI (normalised to its volume) in Cav-1 Null MEFs at 1.0 mg/ml and 1.5 mg/ml concentrations at 15 min (n=58,48, N=6) and 4 hours (n=42,47, N=5). **(G)** Representative XY cross-section images show Cav-1 null MEFs transfected or untransfected with Cav-1 mCherry (magenta) embedded in 1.0 mg/ml and 1.5 mg/ml collagen gels labelled with Phalloidin (green). Graphs show collagen branch junctions and total branch number surrounding the Cav-1 Null reconstituted (top) (n=40,28, N=4) and untransfected Cav-1 Null MEFs (bottom) (n=22,21, N=3) in both collagen concentrations at 4 hours. All graphs show all data points with median and quarters, and error bars mapping spread of data points. Scale bar: 5 μm. Mann Whitney U test for Stiffness studies (C), One-way ANOVA for the rheology (D) and Unpaired T-test have been used for the rest of the graphs. (*P<0.01, **P<0.001, ***P<0.0001, ****P<0.00001)

While these measurements are made on cells adherent and spread on glass with strong focal adhesions, cells in 3D matrices lack strong adhesions (38) and vary in their cell shape and actin organisation (39). Unfortunately, we are unable to make AFM measurements in 3D gels. We plated cells on 2D glass coverslips coated with poly-L-Lysine (PLL), to significantly reduce their adhesion and spreading. Cells were attached mainly but round on PLL. Measured differences in the inherent stiffness of these cells showed cells to be much softer than when spread on glass **(Fig 3B)**. Interestingly, the differential stiffness between WT MEFs and Cav-1 Null MEFs was retained on PLL-attached round cells. This suggests that in a low-adhesion setting, like 3D microenvironment, the difference in the stiffness between these two cell types could be retained. This could contribute to their behaviour at the cell-matrix interphase in 3D gels.

As a first step to understanding the possible effect differential mechanoresponsiveness of WT or Cav-1 Null MEFs could have on 3D collagen gels, we first tested the impact they have on the overall stiffness of the gel. The stiffness of collagen gels depends on fibre thickness, density, and porosity, which rely on its concentration and cross-linking (40,41). Using a parallel plate rheometer, we measured the stiffness of collagen gel (1.0 vs 1.5 mg/ml) at 15 min and 4-hour time points. The storage moduli, *G*′, of unlabelled collagen gels at 37°C measured by rheology, is found to increase with collagen concentration (*G*′ ∼ *c*^2.1^ at 37°C)(40). Our measurements also show the G’ for 1.5 mg/ml collagen gels to be significantly higher than 1.0 mg/ml at 15 min and 4 hours **(Fig 3D)**. This reflects the change in collagen branch numbers and junctions that were seen earlier **(Fig 1D).**

Effect WT and Cav-1 Null MEFs (1 × 10^5^ cells) added to 400 ul collagen (as used earlier to evaluate cell-matrix interaction) have on the G’ was measured. In the presence of either WT MEFs or Cav-1 Null MEFs, G’ for 1.5 mg/ml, collagen gels continue to be significantly higher than 1.0 mg/ml at both time points **(Fig 3D)**, suggesting no significant change in the global stiffness of the gels in the presence of either cell. This is true for this particular cell density and could vary as that changes. Additionally, the viscoelasticity of the gels was looked at by comparing the viscosity (G”) and the elastic (G’) properties of the gels. This was also unaffected by the presence or absence of cells at early or late time points (Sup Fig 2C). This data suggests that the gel’s overall stiffness isn’t affected by this density of WT and Cav-1 Null MEFs **(Fig 2C)**.

3D collagen organisation in 1.0 and 1.5 mg/ml gels with embedded Cav-1 Null MEFs were analysed in the 50-pixel ROI surrounding the cell at 15 min (early) and 4 hours (late) time points **(Fig 3D)**. Like WT MEFs, a collagen ‘cortex’ is seen at the immediate perimeter of the Cav-1 Null MEFs **(Sup Fig 2D)**. When analysed, the number of branches and junctions around the cells significantly increased at both time points. There is no significant difference between the two collagen concentrations at 15 min, comparable to WT MEFs **(Fig 3E)**. However, at 4 hours, unlike in WT MEFs where a significant difference in branch number and junctions between the two concentrations (1.0 mg/ml > 1.5 mg/ml) was seen, no change was detected near Cav-1 null MEFs **(Fig 3E)**. To confirm this is indeed due to a lack of Caveolin-1, Cav-1 Null MEFs were reconstituted with WT Cav-1. At 4 hours, collagen organisation around these transfected cells were comparable to WT MEFs **(Fig 3F**, **Fig 2C)**. Untransfected cells in the same gels behaved like Cav-1 Null MEFs **(Fig 3F**, **Fig 3E)**. This confirms the presence of caveolin-1 and its loss (and the resulting change in cell mechanoresponsiveness) to particularly affect 3D collagen gel organisation at a late time point.

### Actin cytoskeleton regulates cell-dependent collagen organisation

The actin cytoskeleton, downstream of integrins, is a crucial link between the cell and the extracellular matrix (39). Cell stiffness is primarily governed by actin, allowing it to generate the necessary forces to manipulate its surroundings (42). The shape and mechanical properties of the cell are defined by the actin cytoskeleton organised into higher-order arrays capable of dynamic remodelling (43). The actin cortex is also closely associated with the plasma membrane regulating cell mechanics (43–45).

Disrupting the actin cytoskeleton could affect cell stiffness, protrusivity, interaction with and remodelling of the matrix (46,47). Latrunculin A (Lat A), promotes actin filament depolymerisation and sequestering of actin monomers (48,49). Treatment of cells with increasing Lat A concentration for 2 hours causes a concentration-dependent disruption of the actin organisation and a loss in cell morphology in 2D adherent WT and Cav-1 Null MEFs **(Fig 4A)**. This also causes a significant concentration-dependent drop in the stiffness of both WT and Cav-1 Null MEFs, measured using AFM on 2D adherent cells **(Fig 4A)**.

**Figure 4:**
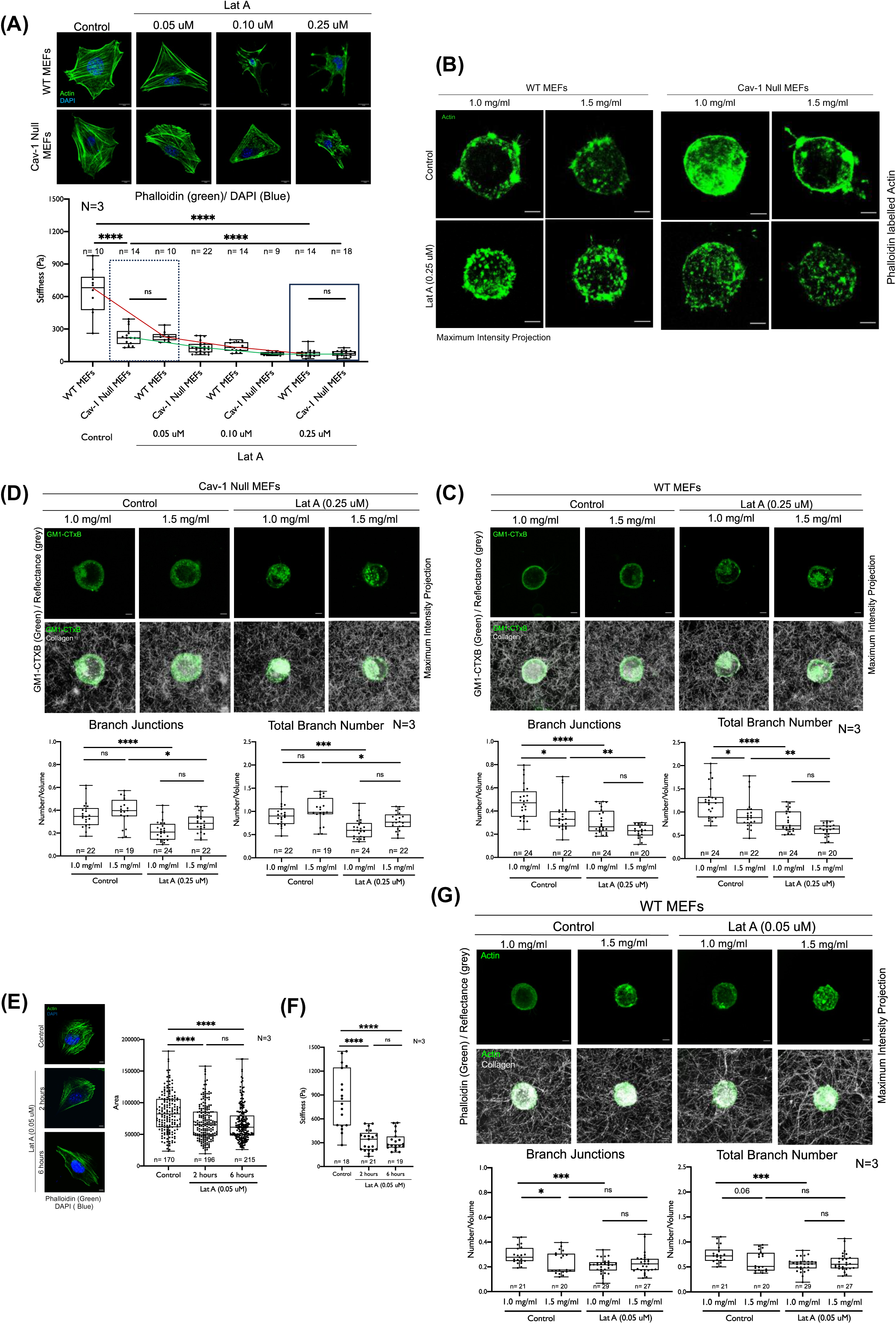
Regulation of collagen organisation near WT MEFs and Cav-1 Null MEFs is actin-dependent: **(A)** Representative images of WT MEFs and Cav-1 Null MEFs, DMSO (control) and Latranculin A (LatA) treated (0.05 μM, 0.10 μM and 0.25 μM Lat A) stained with Phalloidin (green) and DAPI (blue). Graph shows stiffness (Pa) of WT MEFs and Cav-1 Null MEFs, DMSO (Control) and Lat A (0.05 μM, 0.10 μM and 0.25 μM) treated (n=9-18, N=3). The line drawn marks the change in median stiffness across treatments for WTMEFs (red) and Cav-1 Null MEFs (green). **(B)** Representative Z stack MIP image of DMSO (control) and Lat A treated (0.25 μM) WT MEFs and Cav-1 Null MEFs in 1.0 mg/ml and 1.5 mg/ml collagen gels labelled with Phalloidin (green). **(C, D)** Representative Z stack MIP of **(C)** WT MEFs and **(D)** Cav-1 Null MEFs treated with DMSO (Control) or Lat A (0.25 μM) in 1.0 mg/ml and 1.5 mg/ml 3D collagen gels. Cells labelled with GM1-CTxB (green), and collagen imaged by reflectance (grey). Graphs show branch junctions and total branch number (normalised to the volume of ROI) near DMSO (control) or Lat A (0.25 μM) treated WT MEFs (n=24,22,24 and 20, N=3) and Cav-1 Null MEFs (n=22,19,24 and 22, N=3). **(E)** Spreading of WT MEFs on glass treated with Lat A (0.05 uM) for 2 hours and 6 hours (n=170,196 and 215, N=3). The left panel shows representative images of these cells labelled with phalloidin (green) and DAPI (blue). **(F)** Stiffness of WT MEFs treated with DMSO, 0.05 uM Lat A for 2 hours and 6 hours (n=18,21 and 19, N=3 **(G)** Representative Z stack MIP of DMSO (Control) or Lat A treated (0.05 μM) WT MEFs in 1.0 mg/ml and 1.5 mg/ml collagen gels. Cells are labelled with phalloidin (green), and collagen imaged by reflectance (grey). Graphs show branch junctions and total branch number (normalised to the volume of ROI) near cells (n=21,20,29 and 27, N=3). All graphs show all data points with median and quarters and error bars map the spread of data points. Scale bar: 5 μm. Statistical analysis: One way ANOVA. (*P<0.01, ***P<0.0001, ****P<0.00001)

Lat A treatment at 0.25uM, seen to cause a complete disruption of the actin cytoskeleton and cell stiffness **(Fig 4A)**, was chosen for 3D experiments. Lat A treated WT and Cav-1 Null MEFs in 1.0 and 1.5 mg/ml gels, when labelled with phalloidin, showed an expected disruption in actin organisation **(Fig 4B)**. This, however, made defining the cell boundary in treated cells in 3D gels using actin staining difficult. As mentioned earlier, we had seen the GM1-CTxB labelled cell membrane overlap closely with Actin in control cells. Despite their endocytosis, CTxB-bound GM1 is mainly present on the cell membrane **(Fig 4C, 4D, Sup Fig 1A, 3A)**. WT and Cav-1 Null MEFs treated with Lat A for 2 hours before embedding and incubated for 4 hours with Lat A in 3D collagen, showed a significant reduction in collagen branch junctions and numbers at both 1.0 and 1.5 mg/ml concentration **(Fig 4C, 4D)**. This reduction is more prominent in 1.0 than 1.5 mg/ml (Fig 4C). This also affects the organisation of the collagen ‘cortex’ at the cell-matrix interphase (Sup Fig 3B). The increased collagen branching observed in control WT and Cav-1 Null MEFs (Sup Fig 3B) is prominently lost on Lat A treatment.

The difference in collagen branch and junction numbers in 1.0 vs 1.5 mg/ml collagen near WT MEFs **(Fig 2B**, **Fig 4C)** is lost on Lat A treatment **(Fig 4C)**. Near Cav-1 Null MEFs branch and junction numbers are comparable between 1.0 vs 1.5 mg/ml collagen **(Fig 3B**, **Fig 4D),** which remain unaffected by Lat A treatment **(Fig 4D)**. This suggests that the organisation of 3D collagen near cells depends on actin-mediated regulation of cell stiffness and cell mechnoresponsiveness. This also influences how WT MEFs respond to changing collagen concentration (1.0 vs 1.5 mg/ml).

To establish the role of cell stiffness, we asked if we reduced the stiffness of WT MEF and made it comparable to Cav-1 Null MEFs does this affect their behaviour in 3D gels? WT MEFs treated for 2 and 6 hours with a lower 0.05uM concentration of Lat A marginally affected the actin cytoskeleton (confirmed by phalloidin staining) and comparably affected cell spreading **(Fig 4E)**. The stiffness of these cells expectedly dropped within 2 hours of treatment and stayed comparable for 6 hours **(Fig 4F)**. This shows that 0.05uM Lat A treatment for 2 hours reduced the stiffness of WT MEFs, similar to untreated Cav-1 Null MEFs **(Fig 1A, dotted box)**. This drop stays for 6 hours when the experiment ends. Cells are fixed, and collagen organisation is then measured **(Fig 4F)**. This Lat A (0.05uM) treatment of WT MEFs in 3D collagen gels (1.0 and 1.5 mg/ml), causes the difference in branch junctions and numbers seen near the cell to now be lost **(Fig 4G)**. They now resemble the behaviour of Cav-1 Null MEFs, whose stiffnesses they now match **(Fig 3E**, **Fig 4G)**.

These results strongly suggest actin cytoskeleton-dependent cell stiffness to be vital to the organisation of 3D collagen at the cell-matrix interphase. This is further affected by collagen concentration for WT MEFs, but not the softer Cav-1 Null MEFs. This could also be mediated by differences in matrix-dependent Rho signalling (50) and endocytosis (51) in WT vs Cav-1 Null MEFs that we tested for.

### Role of Rho-ROCK signaling in cell-dependent collagen organization

The small GTPase Rho and its effectors play a critical role in actin polymerisation and stability, regulated by mechanical stimuli (52–54). Rho through its downstream effector Rho kinase (ROCK) regulates myosin II and cell contractility (55). This could affect how cells interact with their immediate microenvironment. Migration of cells in 3D environments is regulated by a switch between actin-based protrusions and contraction-driven blebs, regulated by spatial and temporal activation of Rho GTPases (56,57). In 3D matrices Rac activity was less than in cells on 2D matrices (58), though such a regulation of Rho remains unclear. Cav-1 promotes Rho activity by regulating the localisation of Rho inhibitor p190RHOGAP (13). Loss of Cav-1 causes a reduction in basal Rho activation in Cav-1 Null MEFs (9,59,60). This could contribute to the observed differences in 3D collagen organisation near WT vs Cav-1 Null MEFs. Rho contractility in 3D hydrogels is seen to reorganise and align matrix fibres (56,61), which may locally affect matrix organisation.

Targeting Rho-ROCK signalling using Y27632 (20uM) affects actin polymerization and stress fibre formation affecting cell adhesion and spreading(53,62) in both WT and Cav-1 Null MEFs **(Fig 5A).** ROCK inhibition, significantly reduces the stiffness of 2D adherent cells **(Fig 5B)** making them comparable **(Fig 5B)**. This is mediated by a bigger drop in WT MEF stiffness as compared to Cav-1 Null MEFs **(Fig 5B)**.

**Figure 5:**
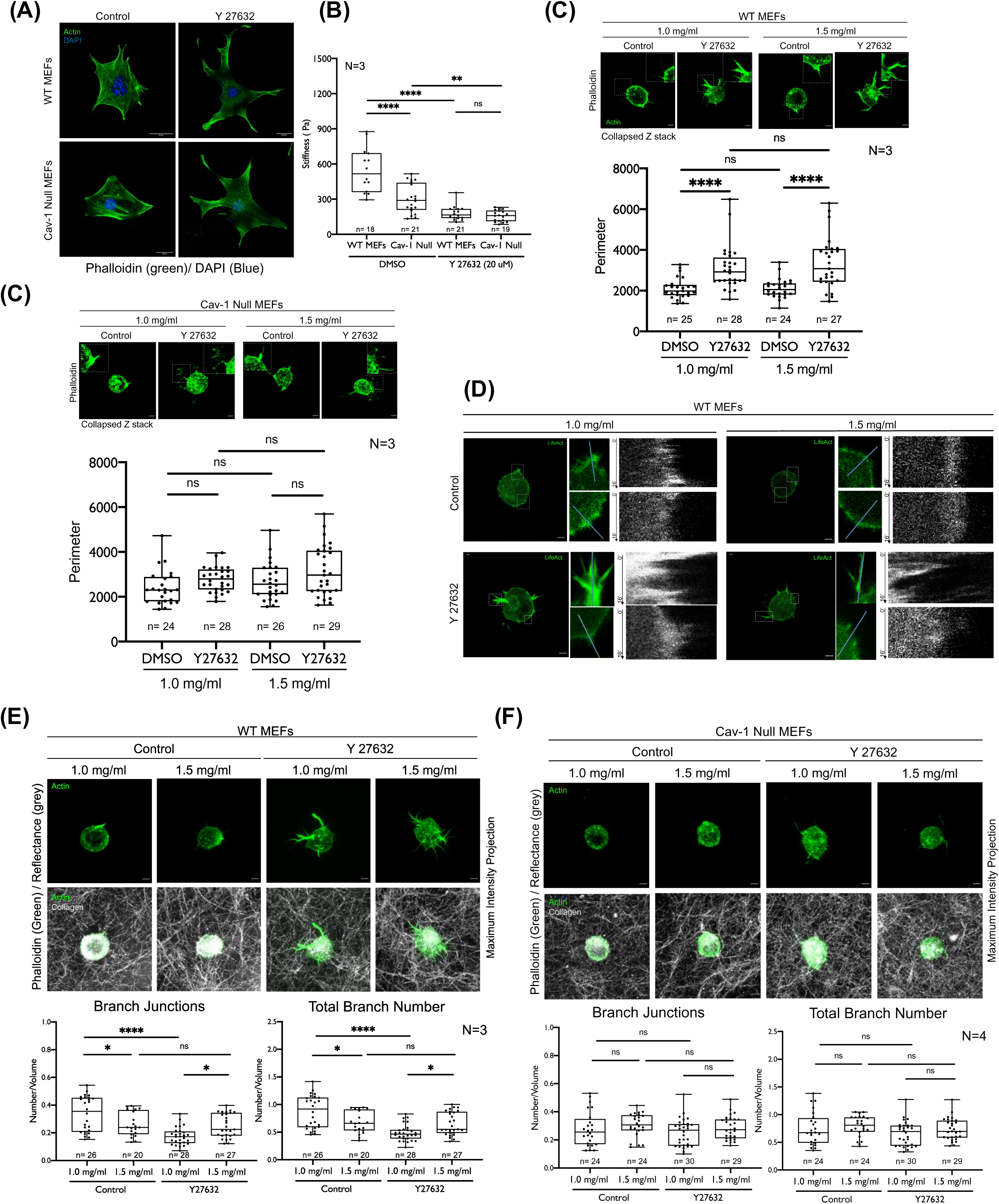
The regulation of collagen organisation near WT MEFs and Cav-1 Null MEFs is dependent on the Rho-ROCK pathway: **(A)** Representative cross-section images of DMSO (control) or Y27632 (20 μM) treated WT MEFs and Cav-1 Null MEFs on glass stained with Phalloidin (green), and DAPI (blue). **(B)** Graphs represent stiffness (Pa) of WT MEFs and Cav-1 Null MEFs on DMSO (control) or Y 27632 (20 μM) treatment measured using AFM (n=18,21,21 and 19, N=3). **(C)** The representative MIP of WT MEFs and Cav-1 Null MEFs treated with DMSO (control) or Y 27632 in 1.0 mg/ml and 1.5 mg/ml 3D collagen gels, stained with Phalloidin (green). Cell protrusions in marked box are zoomed. Graphs show the perimeter of the control or Y 27632 treated WT MEFs (n=25,28,24 and 27, N=3) and Cav-1 Null MEFs (n=24,28,26 and 29, N=3) in 1.0 mg/ml and 1.5 mg/ml 3D collagen gels calculated from the Z stacks. **(D)** Kymographs represent change in fluorescence along a marked line through the membrane in DMSO (Control) and protrusions in Y27632 treated WTMEFs transfected with LifeAct (green) in both collagen concentrations. Representative images of cells with the line marked by a box, zoomed image of the box and kymograph shown. The arrows mark time for the kymograph (0min to 16min). **(E,F)** Representative Z stack MIP of **(E)** WT MEFs and **(E)** Cav-1 Null MEFs treated with DMSO (control) or Y 27632 in 1.0 mg/ml and 1.5 mg/ml 3D collagen gels, labelled with Phalloidin (green), and imaged by reflectance (grey). Graphs represent collagen branch junctions and the number (normalised to the volume of ROI) of 1.0 mg/ml and 1.5 mg/ml collagen gels near DMSO (control) or Y 27632 treated WT MEFs (n=26,20,28 and 27, N=3) and Cav-1 Null MEFs (n=24,24,30 and 29, N=4). All graphs show all data points with median and quarters and error bars map the spread of data points. Scale bar: 5 μm. Statistical analysis: One way ANOVA. (*P<0.01, **P<0.001, ****P<0.00001)

Interestingly, Y27632 treatment of 3D embedded WT MEFs, causes them to make distinct long protrusions in both 1.0 mg/ml and 1.5 mg/ml collagen gels **(Fig 5C)**. This is confirmed by quantitative measurement of their cell perimeter **(Fig 5C)**. Cav-1 Null MEFs, on ROCK inhibition show no significant effect on protrusivity **(Fig 5C)**. Reconstituting Cav-1 Null MEFs with Cav-1 causes increased protrusions to be observed on Y27632 treatment **(Sup Fig 4A)**. This confirms the presence of Cav-1 to be vital to the effect ROCK inhibition has on 3D cell protrusivity **(Sup Fig 4A)**. In WT MEFs embedded in 1.0 and 1.5 mg/ml collagen and treated with Y27632, immunostaining of Cav-1 detected its localization at the membrane but showed no visible enrichment at sites of protrusions **(Sup Fig 4B)**.

Live imaging of LifeAct-GFP transfected WT MEFs and Cav-1 Null MEFs treated with Y27632 shows their protrusivity to be dynamic and impact the collagen matrix in the 50 pixel region surrounding cells over ∼16 min **(Sup Video Fig 1, Sup Fig 4D).** Kymographs for line plotted through protrusions imaged for 50 frames at 11 sec intervals reveal their dynamic nature to be conserved in 1.0 mg/ml and 1.5 mg/ml collagen **(Fig 5D)**. Line plots through membranes that lack protrusions in control or Y27632 treated cells were clearly less dynamic **(Fig 5D)**. This protrusivity doesn’t affect the motility of WT or Cav-1 Null MEFs in the time period they were imaged (16 min) but could over longer periods of time. Videos show the presence of cell protrusions affect the collagen fibre organisation in the 50-pix region around WT MEFs. Lack of protrusion means no such effect is seen in control and Y27632 treated Cav-1 null MEFs **(Sup Video Fig 1, Sup Fig 4C)**. In WT MEFs the Y27632 mediated protrusions while comparable **(Fig 5C)** cause 3D collagen near the cell at 1.0 and 1.5 mg/ml to be affected differently **(Fig 5E)**. Branch junctions and numbers are both seen to decrease significantly in 1.0 mg/ml but largely unaffected at 1.5 mg/ml at 4 hour **(Fig 5E)**. This suggests, small differences in concentration and stiffness of 3D collagen could contribute to their differential behaviour at the cell-matrix interphase. Cell stiffness and protrusivity, both regulated by Rho-ROCK inhibition, could affect these outcomes.

Y27632 mediated ROCK inhibition affects Cav-1 Null MEFs stiffness (making them comparable to inhibitor treated WT MEFs) **(Fig 5B)** but has no visible effect on cell protrusivity **(Fig 5C)**. This interestingly produces no significant change in 3D collagen organisation in proximity to these cells **(Fig 5F)**. It suggests that the drop in stiffness is not enough to affect the cell-matrix interphase.

These findings point to a 3D collagen concentration-dependent response in WT MEFs that is affected by Y27632 treatment, underscoring the relationship between cytoskeletal dynamics and cell stiffness in regulating cell-matrix interphase. Loss of Cav-1 mediated drop in Rho activity(59), should ideally promote protrusivity in these cells, like ROCK inhibition does in WT MEFs. The absence of this suggests there could be additional player(s) that contribute to ROCK-dependent regulation of cell protrusion in 3D collagen gels.

### Dynamin-dependent endocytosis regulates ROCK-dependent protrusivity in WT MEFs

Differential endocytosis in WT vs Cav-1 Null MEFs is known to contribute to differences in signalling and function, regulated among others by cell-matrix adhesion (51). The small GTPase Dynamin is essential for various endocytic pathways, particularly for its role in severing the endocytic vesicle from the plasma membrane through GTP hydrolysis(63–65). Along with its primary role in endocytosis, Dynamin is also implicated in the regulation of actin dynamics (66,67). During endocytosis as well this regulation of actin polymerization is vital. To test the role, Dynamin in ROCK inhibition-dependent protrusivity and regulation of 3D collagen organization in WT MEFs, we used the non-competitive dynamin inhibitor Dynasore. It is known to the GTPase activity of all three Dynamin isoforms (Dynamin 1, 2, and 3) (68).

WT MEFs were pre-treated with only Dynasore for two hours before embedding them in 3D collagen gels and incubated for four hours with the inhibitor. No effect on cell protrusivity was seen **(Fig 6A)**. However, Dynasore when similarly added to cells with Y27632 completely disrupted the protrusivity seen **(Fig 6A**, **Fig 5C)**. This was confirmed by quantitative cell perimeter measurements **(Fig 6A)**. This phenotype was further confirmed using GFP-tagged dominant-negative Dynamin-2 (GFP-Dyn K44A). WT MEFs transfected with Dyn K44A when treated with Y27632 also did not make protrusions in 1.0 mg/ml and 1.5 mg/ml collagen gels **(Fig 6B)**. This says functional Dynamin is needed in WT MEFs for the ROCK inhibition-dependent cellular protrusivity in 3D gels. It is interesting to note that Cav-1 Null MEFs in 3D collagen when treated with Y27632 also make no protrusions **(Fig 5C).**

**Figure 6:**
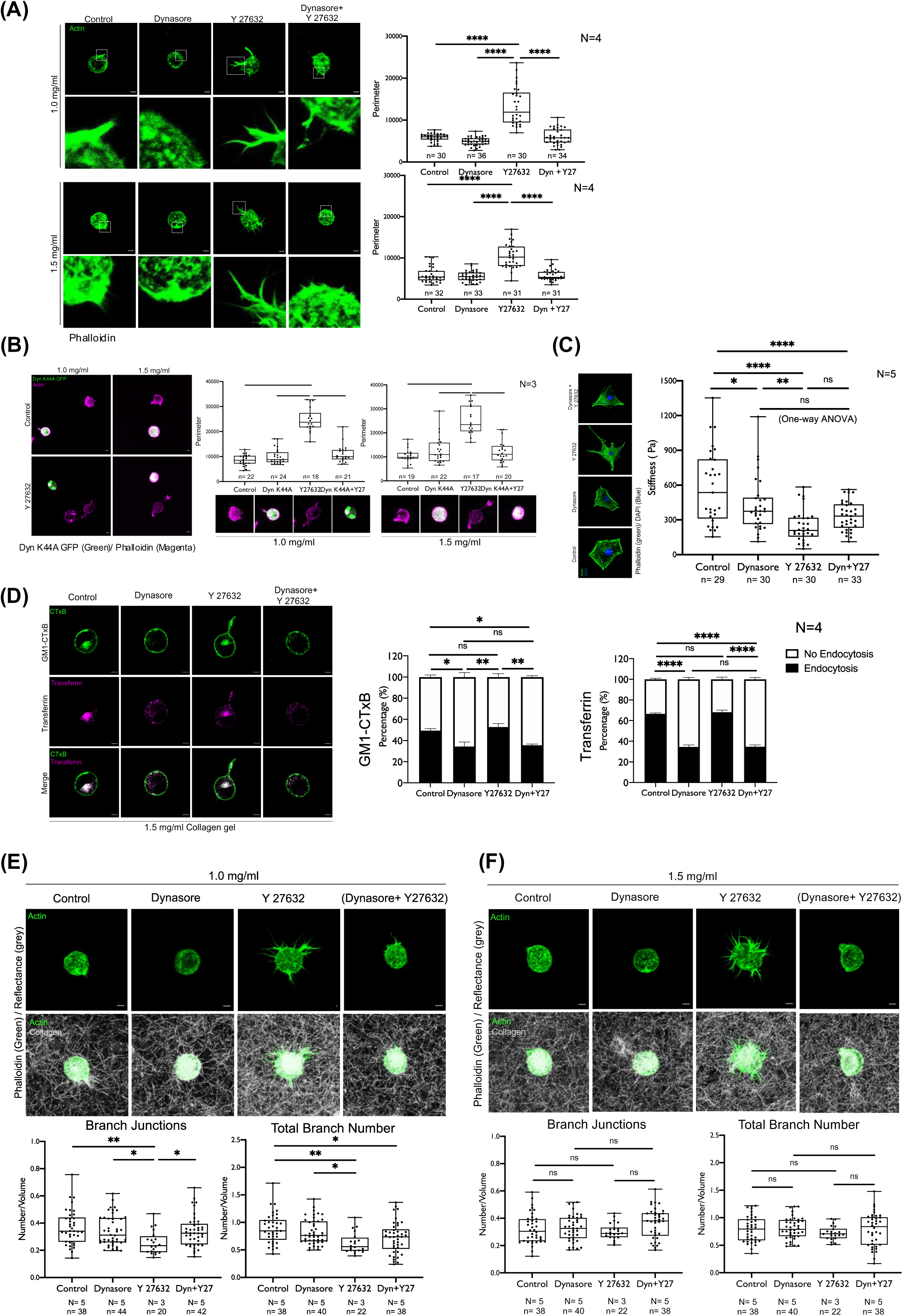
Dynamin regulates ROCK inhibition-dependent effects in the WT MEFs in 3D microenvironment: **(A)** Representative Z stack MIP of WT MEFs in 1.0 mg/ml and 1.5 mg/ml 3D collagen gels treated with DMSO (control), Dynasore, Y27632 and both. Cell protrusions marked by a box shown as zoomed images. The graphs show the perimeter of the cells measured from Z stack images of DMSO (control), Dynasore, Y27632 and Dynasore+Y 27632 (Dyn+Y27) in 1.0 mg/ml (n=30,36,30 and 34, N=4) and 1.5 mg/ml (n=32,33,31 and 31, N=4) 3D collagen gels. **(B)** Representative Z stack MIP of Dyn K44A GFP transfected (or untransfected) WT MEFs (green) and labelled with Phalloidin (magenta) treated with DMSO (control) or Y27632 in 1.0 mg/ml and 1.5 mg/ml 3D collagen gels. Graphs show perimeter calculated from Z stack images of control (untransfected cells with DMSO), Dyn K44A (Dyn K44A GFP transfected cells with DMSO), Y 27632 (Untransfected cells with Y 27632) and Dyn K44A+Y27 (Dyn K44A GFP transfected cells with Y 27632) in 1.0 mg/ml (n=22,24,18 and 21, N=3) and 1.5 mg/ml (n=19,22,17 and 20, N=3) 3D collagen gels. **(C)** Representative cross-section images of WT MEFs on 2D glass coverslips treated with DMSO (control), Dynasore, Y 27632 and Dynasore+Y 27632 (Dyn+Y27), labelled with Phalloidin (green) and DAPI (blue). Graph shows the cell stiffness (Pa) (n=29,30,20 and 33, N=5). **(D)** Representative cross-section images of WT MEFs labelled with CTxB (GM1-CTxB) (green) and Transferrin (magenta) in 1.5 mg/ml 3D collagen gel. Cells were treated with DMSO (control), Dynasore, Y 27632 and Dynasore+Y 27632 (Dyn+Y27) treatment. The graphs show the percentage distribution profile of cells with visible endocytosis (black) or no-endocytosis (white). Graph shows percentage data for >100 cells counted in four individual experiments (N). **(E, F)** Representative Z stack MIP of WT MEFs treated with DMSO (control), Dynasore, Y27632 and Dynasore+Y27632 in (E) 1.0 mg/ml and (F) 1.5mg/ml 3D collagen gels labelled with Phalloidin (green) and reflectance (gray). The graphs show branch junctions and total branch number of **(E)** 1.0 mg/ml and (F) 1.5 mg/ml collagen gels (normalised to the volume of ROI). (E) 38, 44 and 42 images (n) for control, Dynasore and Dyn+Y27 respectively from 5 independent experiments (N) and 20 images (n) for Y 27632 from 3 independent experiments (N) and (F) 38, 40 and 38 images (n) for control, Dynasore and Dyn+Y27 respectively from 5 independent experiments (N) and 22 images (n) for Y 27632 from 3 independent experiments (N) has been analysed. Graphs show all data points with median and quarters and error bars map the spread of data points. Scale bar: 5 μm. Statistical analysis: One way ANOVA (*P<0.01, **P<0.001, ****P<0.00001).

Knowing ROCK inhibition causes a distinct reduction in cell stiffness **(Fig 5A)**, we asked if presence of Dynasore affects cell stiffness too. AFM measurement of 2D adherent WT MEFs showed that Dynasore caused only a small but significant change in cell stiffness **(Fig 6C)**. Dynasore treatment did not significantly alter ROCK inhibition mediated loss of cell stiffness **(Fig 6C)**. Changes in cell stiffness are unlikely to cause the protrusivity changes observed in WT MEFs on Dynasore+Y27632 treatments.

We tested the effect of Dynamin inhibition has on transferrin receptor endocytosis (through clathrin and dynamin-dependent pathways), and CTxB, which binds GM1 and is internalized via Cav-1-dependent (51) or CLIC/GEEC endocytic pathways. In WT MEFs embedded in 1.5 mg/ml 3D collagen gels, Dynasore significantly reduced endocytosis of both transferrin and GM1-CTxB **(Fig 6D)**. This effect persisted when both Dynamin and ROCK were inhibited together. ROCK inhibition alone had no impact on endocytosis **(Fig 6D)**.

Collagen organisation at the cell-matrix interphase, showed ROCK inhibition to significantly reduce number of junctions and branches near WT MEFs embedded in 1.0 mg/ml (but not 1.5 mg/ml) 3D collagen gels **(Fig 5E)**. Dynasore alone did not affect collagen organisation near WT MEFs **(Fig 6E, F)**. When treated with both Dynasore and Y27632, a drop in branch junction and number seen near WT MEFs in 1.0 mg/ml 3D collagen on Y27632 treatment is reversed **(Fig 6E)** without affecting 1.5 mg/ml 3D collagen gels **(Fig 6F)**.

This supports our earlier observation that small differences in 3D collagen concentration and stiffness could contribute to their differential behaviour at the cell-matrix interphase. Cell stiffness and protrusivity, regulated by Rho-ROCK signaling mediated regulation of actin and dynamin-dependent endocytosis, contribute to these outcomes. This reflects in the altered behaviour of Cav-1 Null MEFs in 3D collagen gels.

## DISCUSSION

In 3D collagen gels, the number of fibres and their junctions, indicative of density and crosslinking, serve as key markers for assessing how cells modulate their surrounding collagen matrix. Cell influence collagen organisation in their microenvironment through various mechanisms, including 1. secretion of ECM proteins, altering both the concentration and cross-linking, 2. secretion of MMPs to degrade ECM proteins and 3. exertion of mechanical forces on the matrix via the actin cytoskeleton (9,22). Our study examines collagen organisation at two slightly different concentrations at 15 min and 4 hours post-polymerization, and how cells can sense and respond to cues from their immediate 3D microenvironment. Studies suggest that, during this time frame, cell-mediated changes in 3D collagen gels are minimally affected by MMP inhibition (31). They also allow for a change in the behaviour of a “colloidal” collagen gel as it settles and cures over time. These events result in measurable changes in fibre density (branch number) and crosslinking (branch junctions), which could affect the function of embedded cells (11)This, however, is not commonly studied and could offer insights into how cell-matrix interphase impacts cell function.

Defining what can be considered as the **‘immediate vicinity’** of cells for such a cell-matrix interphase becomes essential and we have done this by looking at a 50-pixel region (∼3.25 µm), of the 3D collagen gel around the cell, which our earlier studies (23) have seen to be better than 10 or 30 pixels, where the data was highly variable. Consolidating across Z-stacks allows us to build a 3D collagen region of interest (cell-matrix interphase) that we could compare between cells, cell types and treatments.

Various biochemical and biophysical properties of the cell and the ECM, influence the cell-matrix interphase. Malandrino et al. demonstrated that collagen densification (an increase in local fibre concentration and crosslinking) in this interphase occurs through filopodia-mediated ECM compaction and depends on ECM crosslinking (31). While our observations corroborate this, an intriguing question remains: does the gel’s polymerisation properties solely dictate this process, or does the mechanical sensitivity and activity of the cell also play a critical role? In comparing WT and Cav-1 Null MEFs, with variable mechanosensitivity, we find cells embedded in 3D collagen gels cause the collagen to organise at the cell-matrix interphase (50 pixel) in a way that responds to their inherent stiffness and actin-dependent responsiveness (push and pull).

The presence of a distinct, densely crosslinked region of extracellular collagen closer to the cell could reflect the impact cells have on the matrix. The softer Cav-1 Null MEFs supporting lesser branch numbers and junctions (at lower concentration) also confirm cell stiffness’s impact on collagen organisation. WT MEFs made softer by Lat A treatment (at lower concentrations), behave similarly. When the actin cytoskeleton is disrupted, cells lose their inherent stiffness and responsiveness in 3D gels. These drug-treated significantly softer cells do support some collagen organisation in their vicinity, suggesting how, along with their stiffness and cytoskeletal responsiveness, cells, when in 3D gels, could also provide a surface for the nucleation of collagen fibres regulating their organisation.

Cav-1 and the actin cytoskeleton have a dynamic regulatory relationship, influencing each other’s organisation and function (9,74–77), mechanosensitivity of Cav-1 and the mechanoresponsiveness of actin are also linked. An intact actin network is essential for maintaining caveolae homeostasis, disassembly, and endocytosis in response to mechanical cues (8,9,69). Similarly, the mechanoresponsiveness of actin is Cav-1 dependent, as evidenced by its role in actin cortex reassembly on hypoosmotic shock recovery (70). Cav-1 is critical for actin-mediated ECM contraction through Rho-ROCK signalling (71). It promotes Rho-ROCK and force-dependent actomyosin contraction, driving ECM fibrillogenesis and increasing matrix stiffness, significantly impaired in Cav-1 Null MEFs (11). Caveolar endocytosis is also responsive to mechanical stimuli that affect membrane tension, such as loss of adhesion (51). This needs a functional actin cytoskeleton, possibly regulated by Rho-ROCK-dependent signalling.

In WT MEFs, ROCK inhibition enhances cellular protrusivity in 1.0 and 1.5 mg/ml collagen gels, affecting collagen organisation more prominently around cells in 1.0. This protrusivity may be mediated through Rac, inversely regulated by the Rho-ROCK pathway (72). ROCK inhibition is known to impair cytoskeletal contractibility, which could also affect filopodia formation (73). The protrusions seen in WT MEFs on ROCK inhibition are distinctly bigger in size than filopodia whose role in the 3D microenvironments is well-documented (31). They affect collagen organisation around Y27632 treated WT MEFs in 1.0 mg/ml only. The softer and less crosslinked 1.0 mg/ml gels are affected more prominently by WT MEFs, which could be why ROCK inhibition impacts them. Softer Cav-1 Null MEFs with known changes in actin cytoskeletal regulation and endocytosis are unaffected by changing collagen 3D gel concentrations (1.0 vs 1.5 mg/ml) and hence Y27632 treatment. Dynasore-mediated targeting of Dynamin in disrupting ROCK-dependent protrusivity highlights the critical role dynamin-dependent caveolar endocytosis could have in this pathway. Dynasore treatment in 3D embedded cells affecting transferrin and GM1 endocytosis in WTMEFs supports this regulatory crosstalk.

The role dynamin-dependent endocytosis has in supporting this regulation remains speculative at best. Changes in endocytic activity can significantly alter the composition of plasma membranes, affecting mobility in the membrane and its mechanical properties (74,75). This could also influence the recruitment of Rho GTPase and Rho GEFs or GAPs to regulate Rho-ROCK signalling, affecting inhibition-mediated protrusivity and collagen remodelling. Beyond its role in endocytosis, dynamin is reported to regulate actin polymerisation (66), which could also be impacted by Dynasore treatment to disrupt ROCK-inhibition mediated protrusivity. Dynasore only marginally affects cell stiffness, supporting its regulation of endocytosis to be more relevant in this pathway.

In summary, our findings suggest that cells in 3D collagen gels actively promote its accumulation at the cell-matrix interphase through ROCK-mediated, actin-driven contraction. This accumulation is influenced by the degree of collagen crosslinking, with less crosslinked collagen (1.0 mg/ml) more susceptible to cellular remodelling. ROCK activation, dependent on Cav-1, plays a critical role in this process. When ROCK activity is lost, the loss of actin-driven contraction is disrupted, leading to reduced collagen accumulation at the cell matrix interphase. Notably, this effect is more pronounced in 1.0 mg/ml collagen gels, where initial collagen alteration by cells is higher. Interestingly, inhibiting dynamin restores collagen organisation in 1.0 and 1.5 mg/ml gels, suggesting dynamin activity to be essential for actin-dependent regulation of cell behaviour in 3D. Dynamin’s involvement in regulating endocytic pathways could be a major player here. These findings highlight the intricate interplay between biochemical pathways, cellular mechanics, and the extracellular matrix in regulating collagen organisation around cells.

## Author’s Contribution

Conceptualisation: N.B, D.M. Experiments: S.K. D.M. (AFM), G.P. (Rheology), D.M. (everything else), Analysis: S.K. S.P. (AFM), G.P. M.D. (Rheology) D.M. A.K. N.B. (everything else) Original Draft: D.M. N.B. Review and Editing: S.P. M.D. N.B. D.M. Fund acquisition and project supervision: N.B.

## Acknowledgement

This work is supported by SERB CRG grant-CRG/2022/001813 to NB. We thank Prof Richard Anderson for the cell lines WT MEFs and Cav-1 Null MEFs and Prof Aurnab Ghosh for LifeAct GFP construct. We acknowledge the extensive support provided by IISER Pune microscopy facility for imaging.

