## Supplementary material for "Caveolin-1 dependent regulation of cell-matrix interphase in 3D collagen gels": Supp Figures

### Supplementary Figure 1

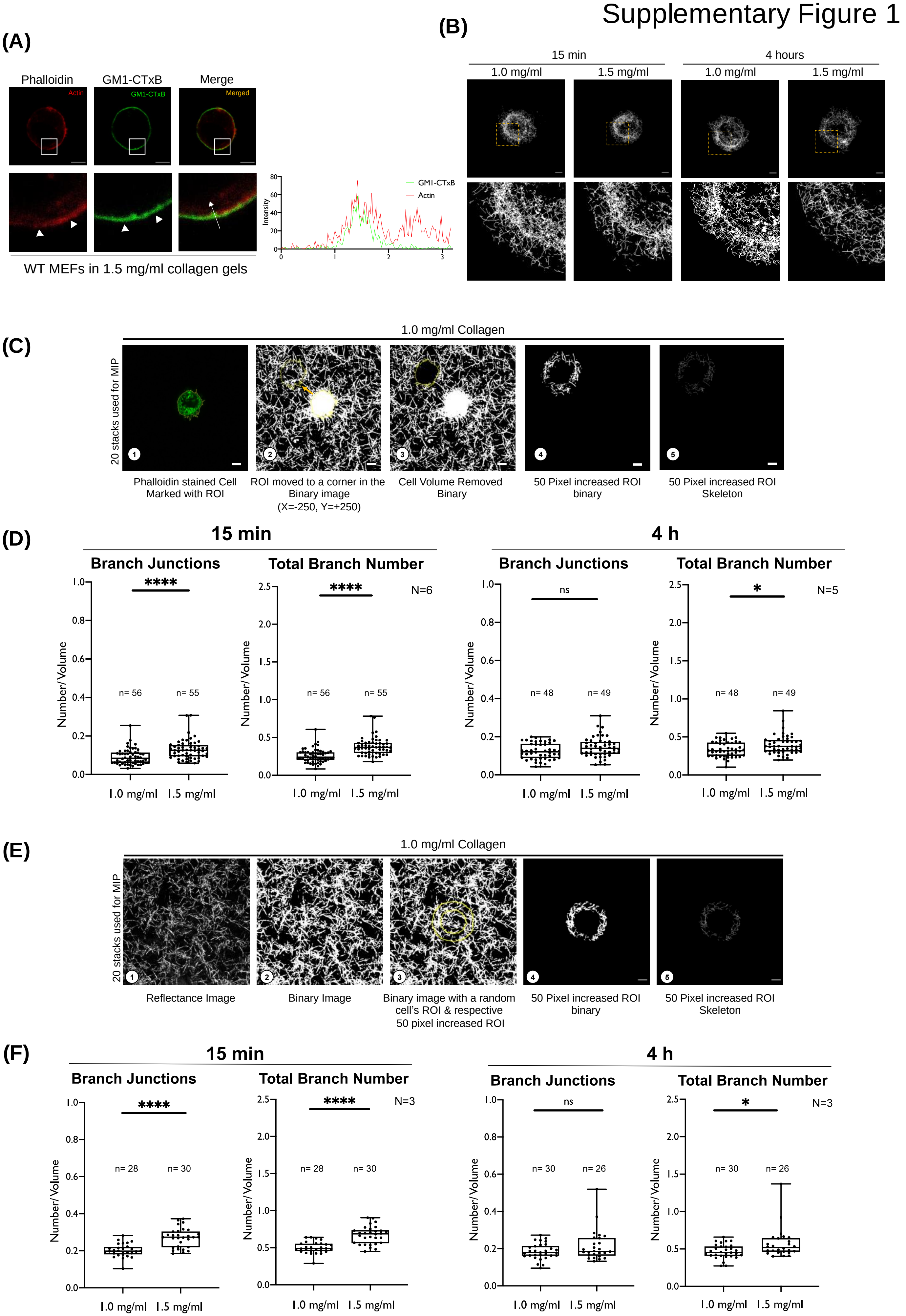

**Supplementary Figure 1: 3D Collagen organisation away from WT MEFs. (A)** Representative images of WT MEFs in 1.5 mg/ml collagen gels labelled with GM1-CTxB (green) and phalloidin (Red). Line plots fluorescence intensity of GM1-CTxB (green) and actin (red) along marked line in the cell. **(B)** The representative Z stack MIP of skeletonised collagen around WT MEFs (50-pixel ROI) in 1.0 mg/ml collagen gels at 15 min and 4 hours (Top panel). Skeletonised collagen marked by a box is zoomed (Bottom panel). Stepwise evaluation of collagen organisation **(C)** away from the cell in 3D gels. A representative MIP of 20 cross-sections from a Z-stack used. (1) Phalloidin-stained WT MEFs (green) marked with ROI (2) pasted on the binary reflectance image, moved to corner of gel image by X=-250, Y-+250 pixels. (3) Signal from inside ROI removed, (4) ROI increased by 50-pixels (Fig 1A), and signal outside removed to get a 50-pixel ROI binary, (5) skeletonised for analysis. **(D)** Graphs show branch junctions and total branch number of collagen in 50-pixel region away from WT MEFs (normalised to ROI volume) in 1.0 mg/ml and 1.5 mg/ml gels at 15 minutes (n=56,55, N=6) and 4 hours (n=48,49, N=5). **(E)** Stepwise workflow for evaluating collagen organisation in 3D gels without cells. (1) Representative reflectance image of 1.0 mg/ml 3D collagen gel (2) converted to binary image (3) 50-pixel ROI pasted, (4) ROI binary generated, (5) skeletonised for analysis. **(F)** Graphs show the branch junctions and total branch number (normalised to ROI volume) in 1.0 mg/ml and 1.5 mg/ml collagen gels at 15 min (n=28,30, N=3) and 4 hours (n=30,26, N=3). All graphs show all data points with median and quarters. Error bars map the spread of data points. Scale bar: 5 μm. Statistical analysis: Unpaired T test (*P<0.01, ****P<0.00001)

Supplementary Figure 2

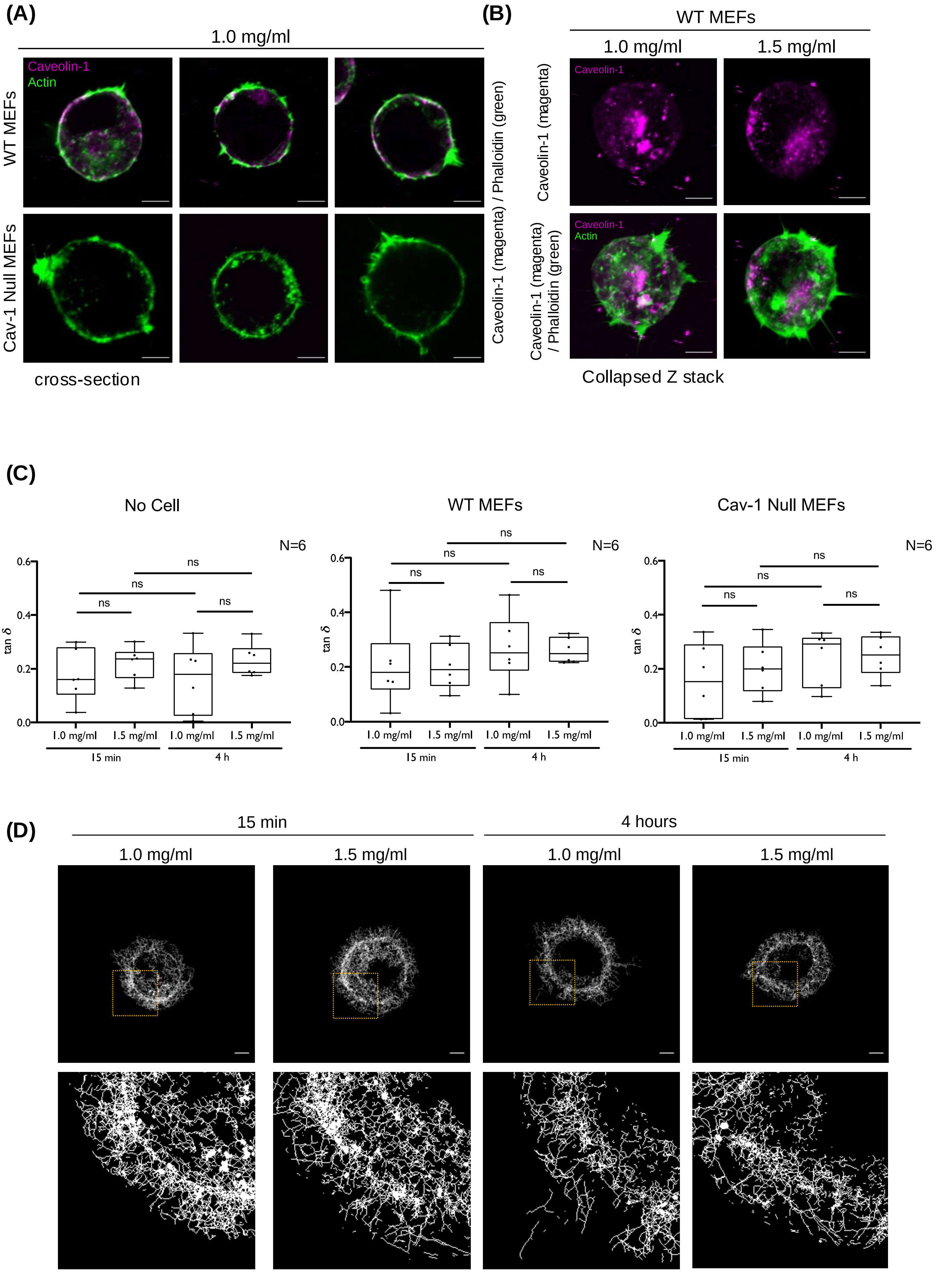

**Supplementary Figure 2: Characterization of Cav-1 Null MEFs and collagen gels. (A)** Representative cross-section images of WT MEFs and Cav-1 Null MEFs in 1.0 mg/ml collagen gels and **(B)** Z stack MIP of WT MEFs in both collagen gels labelled for Caveolin-1 (magenta) and actin (green) **(C)** Tan Delta (tan $\boldsymbol{\delta}$) of 1.0 mg/ml and 1.5 mg/ml collagen gels without cell, WT MEFs and Cav-1 Null MEFs for 15 min and 4 hours (4h) at 0.1 rad/s angular frequency (N=6) **(D)** Representative Z stack MIP of skeletonised collagen around Cav-1 Null MEFs (50 pixel ROI) in 1.0 mg/ml and 1.5 mg/ml 3D collagen gels at 15 min and 4 hours (Top panel). Skeletonised collagen marked by box shown as zoomed image in bottom panel.

Supplementary Figure 3

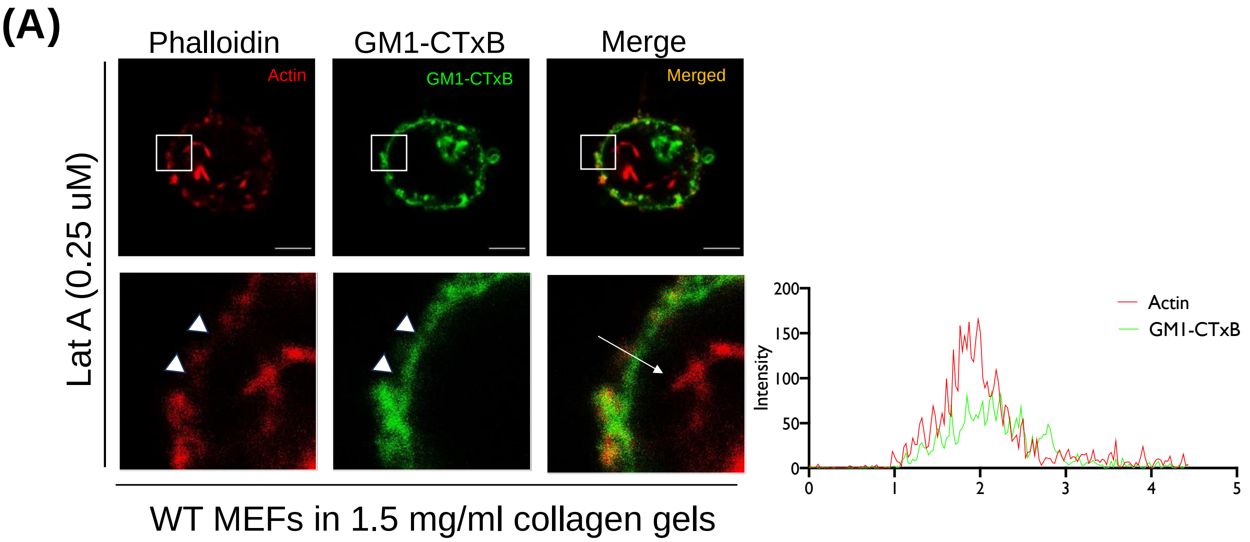


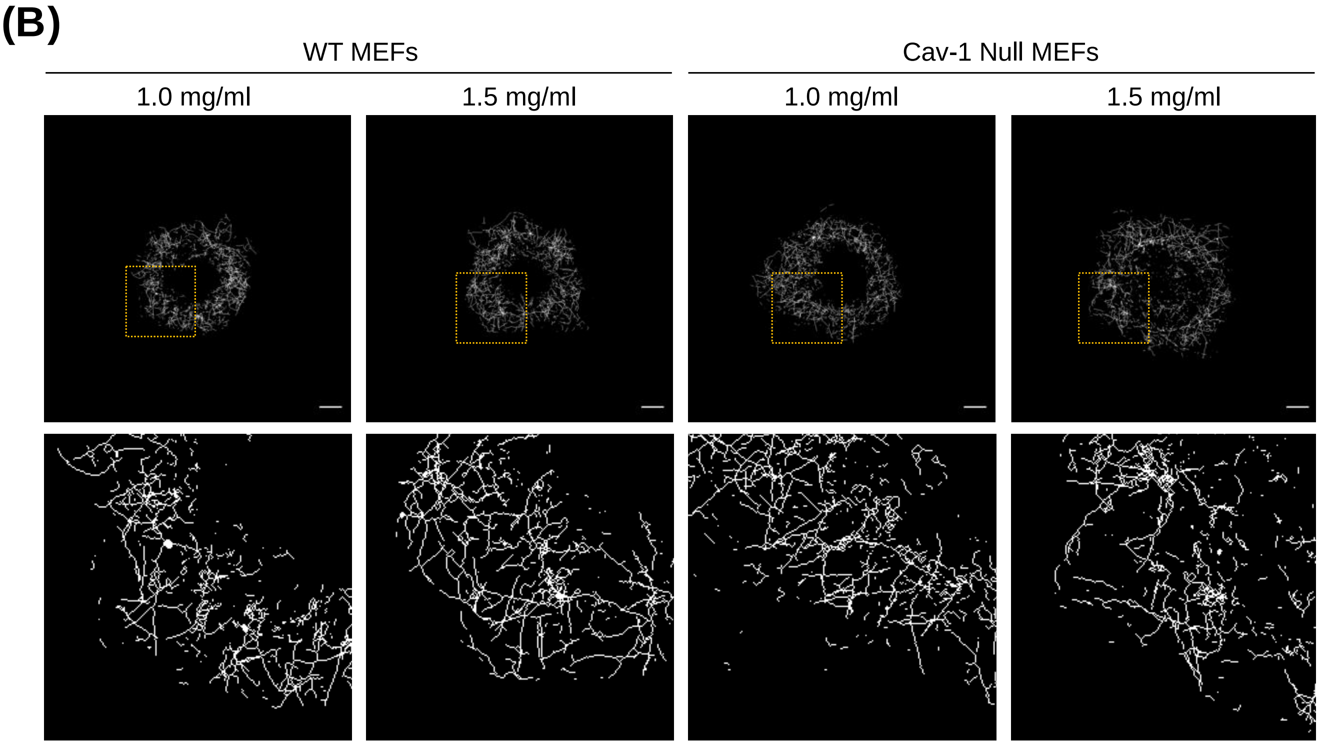

**Supplementary Figure 3: GM1-CTxB labelling to mark cell boundary in Lantranculin A treated cells: (A)** Representative images of WT MEFs in 1.5 mg/ml collagen gels labelled with GM1-CTxB (green) and phalloidin (Red) on Lat A (0.25M) treatment. Line plots fluorescence intensity of GM1-CTxB (green) and actin (red) along marked line in LatA-treated cells. **(B)** The representative Z stack MIP of skeletonised collagen around the WT MEFs and Cav-1 Null MEFs (in a 50-pixel ROI) in 1.0 mg/ml and 1.5 mg/ml 3D collagen gels (Top panel). Skeletonised collagen marked by a box are shown as zoomed in image (Bottom panel).

### Supplementary Figure 4

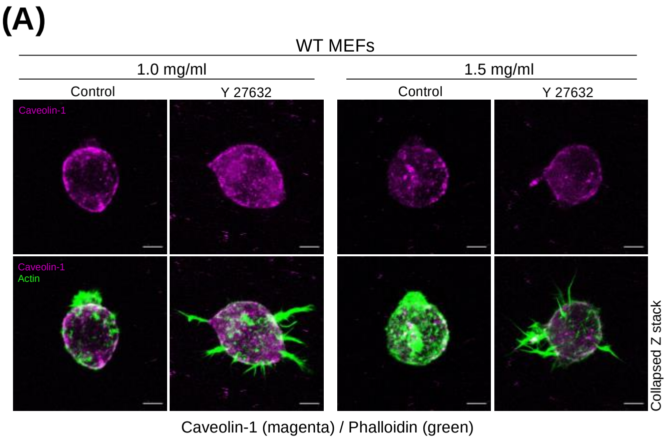


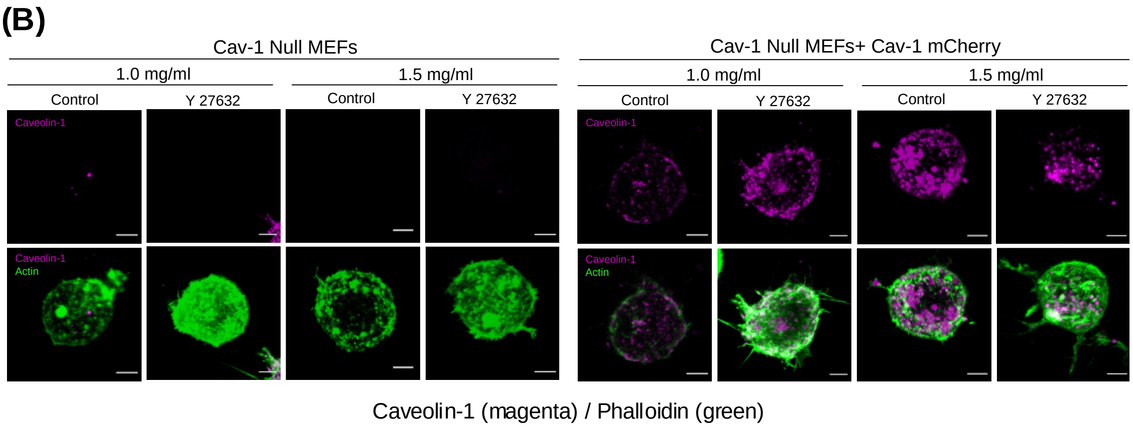


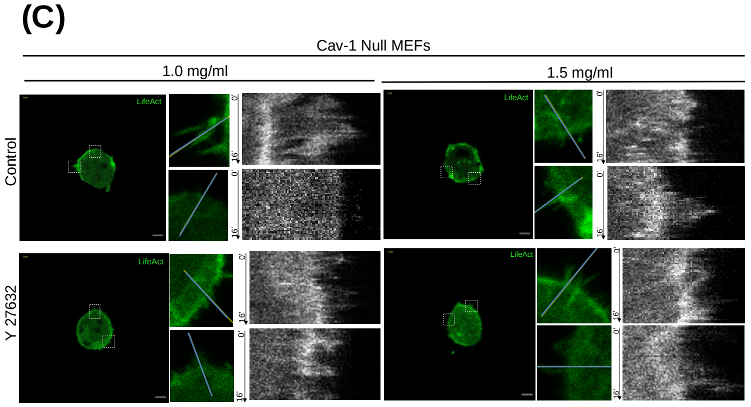


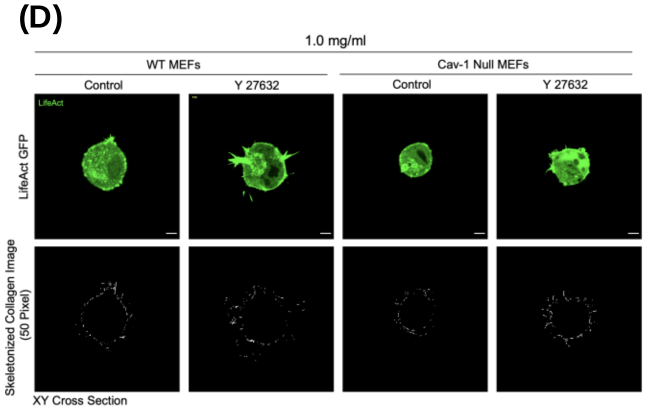

**Supplementary Figure 4: Effect of Rho-ROCK pathway inhibition on WT MEFs and Cav-1 Null MEFs in 3D collagen gels. (A)** Representative Z-stack MIP of DMSO (control) or Y 27632 treated WT MEFs stained with Cav-1 (Magenta) and Phalloidin (green) in 1.0 mg/ml and 1.5 mg/ml 3D collagen gels. **(B)** Representative MIP showing Cav-1 Null MEFs untransfected or transfected with Cav-1 mCherry (magenta) and labelled with Phalloidin (green) treated with DMSO (control) or Y 27632 treated in 1.0 mg/ml and 1.5 mg/ml 3D collagen gels. **(C)** Kymographs represent changes in fluorescence on marked lines through membrane or protrusion in DMSO (Control), Y27632 treated LifeAct GFP transfected Cav-1 Null MEFs (green) in 1.0 mg/ml and 1.5 mg/ml 3D collagen gel. Representative images of cell, line marked by box and zoomed and kymograph shown. Arrow marks time of the kymograph (0min to 16min). **(D)** Cross-section image of one movie frame for DMSO (control), Y 27632 treated WT MEFs and Cav-1 Null MEFs in 1.0 mg/ml collagen gels. Top panel - LifeAct GFP (green) and bottom panel collagen skeletonised image in a 50-pixel ROI near cell. Movies are provided independently. Scale bar: 5 μm.

**Supplementary Video 1: WT MEFs and Cav-1 Null MEFs treated with DMSO (control) or Y 27632 in 1.0 mg/ml collagen gel:** WT MEFs and Cav-1 Null MEFs transfected with LifeAct GFP (green) treated with DMSO (control) or Y 27632. Cell with LifeAct-GFP shown in green and collagen reflectance image shown in grey. Scale bar: 5 μm.

LifeAct GFP expressing WTMEFs in 1.0 mg/ml collagen treated with, **Video 1 and 2:** DMSO (Control) (1) skeletonised collagen in 50-pixel ROI (2) or **Video 3 and 4:** Y 27632 (3) skeletonised collagen in 50-pixel ROI (4).

LifeAct GFP expressing Cav-1 Null MEFs in 1.0 mg/ml collagen treated with, **Video 5 and 6:** DMSO (Control) (5) skeletonised collagen in 50-pixel ROI (6). **Video 7 and 8:** Y 27632 (7) and skeletonised collagen in 50-pixel ROI (8).
